# A simple liquid 3D cell culture paradigm models oxidative mitochondrial metabolism of epithelial breast cancer cells with relevance for lung metastases

**DOI:** 10.1101/2025.08.24.671623

**Authors:** Kuppusamy Balamurugan, Melissa R. Mikolaj, Jonathan M. Weiss, Ronald Holewinski, Yu Fan, Xia Xu, Lois McKennett, Christopher W. Dell, Duncan Donohue, Ariana Vitale, Shashikala Ratnayake, Shikha Sharan, Qingrong Chen, Daoud Meerzaman, Thorkel Andresson, Daniel W. McVicar, Kedar Narayan, Esta Sterneck

## Abstract

Three-dimensional (3D) cell culture systems have emerged as powerful tools for modeling tumor biology in ex vivo settings. However, the diverse array of available 3D culture methods presents challenges in selecting the most appropriate model for specific research questions. This study provides a comparative analysis of breast cancer cells (SUM149, IBC-3, and MDA-MB-468) in the mammosphere culture (SphC) model or an “emboli” culture (EmC) model, which enrich for cancer stem cells and epithelial features, respectively. The EmC model, designed originally for inflammatory breast cancer, is characterized by media viscosity and mechanical rocking of the culture vessel. Notably, cells in EmC showed a distinct and durable reduction in cell proliferation ex vivo while demonstrating increased capacity to establish experimental lung metastases in vivo. Ultrastructural quantitative analysis of electron microscopy images suggested that cells in EmC acquire nuclear and mitochondrial features that resemble those of tumor tissue. Proteomics, single-cell transcriptomics, and metabolic flux analyses showed that cells in EmC and SphC favor mitochondrial oxidative metabolism (OXPHOS) and glycolysis, respectively. EmC rendered cells hypersensitive to OXPHOS inhibition, but more resistant to oxidative stress. Several genes associated with lung metastasis, including ID1, were specifically enriched in EmC. Given the emerging role of OXPHOS in cancer cell survival during dissemination and as established metastases, we propose that the EmC paradigm is a suitable ex vivo model to study signaling pathways relevant for tumor tissue and to assess drug sensitivities and resistance mechanisms of metastatic breast cancer cells ex vivo.

**SIGNIFICANCE:** This study provides an in-depth characterization of a resource-efficient yet powerful 3D culture paradigm to improve the physiological relevance of ex vivo approaches. Applicable to epithelial cancers, this model offers a platform to accelerate the discovery of physiologically relevant signaling pathways and specific cancer cell vulnerabilities.

## INTRODUCTION

For over a century, cell culture has been a workhorse of biological research, conventionally conducted in two dimensions (1,2). Although 2D cultures provide ease of use and reproducibility, they are limited by nonphysiological substrate stiffness and favor rapidly proliferating cell lines. To better recapitulate physiological conditions, a variety of three-dimensional (3D) culture paradigms are being developed with varying degrees of complexity including scaffolds or matrices, various cell types, and/or dynamic features (3–6). Each system presents distinct limitations and advantages depending on the research question to be addressed.

We previously demonstrated that metastatic breast cancer cells cultured in a liquid 3D paradigm, but not in 2D, recapitulate an *in vivo* signaling pathway that preserves E-cadherin protein expression even when its mRNA is downregulated (7). Epithelial-mesenchymal transition (EMT), a well-studied mechanism of metastasis, involves downregulation of genes for epithelial proteins like E-cadherin and induction of mesenchymal proteins such as vimentin. Driven by EMT transcription factors, it promotes stemness and invasiveness (8,9). EMT gene signatures are notably upregulated in metastatic cancer (10) and contribute to the metastatic process (11,12). However, most invasive ductal carcinomas and overt breast cancer metastases retain E-cadherin protein expression, and several studies have reported a positive correlation between E-cadherin, metastasis, and poor prognosis (13–18). Thus, an ex vivo metastatic breast cancer culture model that preserves E-cadherin-mediated cell-cell adhesions may provide a more physiologically relevant platform for studying metastatic biology and therapeutic vulnerabilities.

The 3D paradigm used in our prior study (7), originally developed by Lehman et al. specifically to model intralymphatic emboli by inflammatory breast cancer (IBC) cells, incorporates lymph-like media viscosity and slow rocking for mild mechanical stimulation (19). We extended this emboli culture (EmC) model to additional breast cancer cell types, enabling the study of cell phenotypes under conditions where cell-cell adhesions are maintained in all dimensions. Here, we present a systematic characterization of breast cancer cells cultured in EmC compared with 2D cultures, and in some contexts, with tumor tissue. Furthermore, because it is a widely used liquid 3D model, we included sphere culture (SphC), which was developed specifically to select for cancer stem cell phenotypes (20,21). Three basal epithelial cell lines were analyzed: two IBC cell lines, the triple-negative breast cancer (TNBC) SUM149 cells and HER2+ IBC-3 cells, as well as MDA-MB-468 cells, also representing TNBC. Our results indicate that the 3D EmC promotes many features observed in primary tumors and lung metastases in breast cancer. Given the current urgency to increase research efforts with human ex vivo models, we propose EmC as a resource-efficient, yet powerful ex vivo approach to advance the identification of context-specific signaling pathways and actionable therapeutic vulnerabilities.

## MATERIALS AND METHODS

### Cell culture and cell death assays

SUM149 (RRID:CVCL_3422) and MDA-IBC-3 (RRID:CVCL_HC47) cells originated from Asterand Bioscience and MDACC, respectively. MDA-MB-468 (RRID:CVCL_0419) and MCF-7 (RRID:CVCL_0031) cells were from ATCC. MDA-MB-231-LM2 (RRID: CVCL_5998) and SUM190 RRID: CVCL_3423) cells were a kind gift from Dr. J. Massague (MSKCC) and Dr. P. Steeg (NCI), respectively. Cells were used at passages 2-30, last authenticated in 2025 by GenePrint®10 (Promega), and routinely *Mycoplasma* tested by qPCR. SUM149 expressing luciferase were as described (7). Cells were cultured at 5% CO_2_/37°C in media with 10% FBS, 100 units/ml penicillin and 100 µg/ml streptomycin as follows: MDA-MB-468, MDA-MB-231-LM2 and MCF-7 in Dulbecco’s modified Eagle’s medium (GIBCO DMEM #11965-092), SUM149, SUM190 and IBC-3 in Ham’s F-12 media (GIBCO #31765092) with 5 μg/ml hydrocortisone (MilliporeSigma #H-0135) and 1 μg/ml Insulin (MilliporeSigma #I-0516). In addition, all media contained 2.25% PEG8000 (MilliporeSigma #202452) except for sphere cultures (SphC), which were cultured in Mammocult media (StemCell Technologies #05620) supplemented with 2 μg/ml Heparin (StemCell Technologies #07980) and 0.44 μg/ml hydrocortisone. Emboli culture (EmC) was performed essentially as described (19) except that 6-well or 24-well ultra-low attachment (ULA) dishes were used (Corning #3471, #3473). Briefly, 2.5 x 10^5^ cells were seeded in 6-well plates in 2 mL medium and gently rocked at approximately 40 rpm for 3 days unless otherwise indicated. Fresh medium (1 mL) was added on day 3 for longer culture periods. IACS-10759 (#S8731), sodium dichloroacetate (DCA; #S8615) and AGX51 (#E1152) were from Selleck Chemicals, FX-11 from CAYMAN Chemicals (#213971-34-7), and DMSO from MilliporeSigma (#D-2650).

To isolate the 3D cell assemblies, cells were centrifuged at 500 rpm for 30 sec, washed with PBS, and treated with TrypLExpress (GIBCO #12604-013) for 10 min with intermittent pipetting before neutralization with cell culture medium. Cell numbers were counted by mixing 10 µl of cell suspension with 10 µl of Trypan Blue dye and analyzing by dye exclusion using a Countess II FL (Life Technologies, USA). For cell proliferation measurements, about 2,000 cells were seeded into 96-well F-bottom black microplate (Griener #655090) and cell numbers were measured at the indicated times using a Celigo imaging cytometer (Nexcelom Biosciences). To assess the number of dying cells, cells were treated with 5 μg/ml propidium iodide (PI; MilliporeSigma #P4130) for 30 min and analyzed by PI-based cell death assay using a Celigo imaging cytometer (Nexcelom).

### Flow Cytometry

For cell cycle analysis, cells were cultured and trypsinized as described above. Ice-cold 70% ethanol (∼500 µl) was added slowly while vortexing the tubes and kept at 4°C for 2 hours. Cells were washed once with 1X FACS washing buffer (PBS + 0.1% BSA) and 200 µl of the propidium iodide staining solution (50 µg/ml PI + 3.8 mM citrate buffer + 10 ng/ml RNase A) was added and incubated for overnight at 4°C. Samples were washed with FACS washing buffer and analyzed by flow cytometry.

For measurement of reactive oxygen species (ROS), 50,000 cells were resuspended in 400 µl serum-free medium, followed by incubation with CM-H2DCFDA (5 µM; #6827, ThermoFischer Scientific) for 20 min in the dark at 37°C. After 30 min, 400 µl ice-cold PBS + 0.5% BSA were added to stop the reaction, mixed well, and the cells were pelleted at 1,500 rpm for 5 min at RT. The pellet was resuspended in 400 µl PBS + 0.5% BSA. To the DCF-DA preloaded cells, 100 µM H_2_O_2_ was added and incubated for 15 min at RT in the dark. The cells were then pelleted (1,500 rpm, 5 min), resuspended in 200 µl PBS + 0.5% BSA, and transferred to a 96-well plate (Griener, #655090). The DCF fluorescence was measured with a Spectramax iD3 plate reader (Molecular Devices, USA) using excitation (492 nm) and emission wavelengths (535 nm) settings.

### Seahorse Cell Mito Stress Test

Before the samples were loaded, the seahorse 96-well microplate was coated with 30 µl of Poly-L-Lysine (MilliporeSigma #P4707) and kept inside the hood for 20 min. After the material was decanted, wells were washed twice with cell culture grade water, incubated inside the hood for 30 min, then placed in a non-CO_2_ incubator at 37°C for 30 min. This coated plate was taken for Seahorse Cell Mito Stress Test. The Seahorse XF Cell Mito Stress Test (Cat#103015-100, Seahorse Bioscience, MA, USA) uses modulators of respiration that target components of the electron transport chain in the mitochondria to measure key parameters of metabolic functions. The oxygen consumption rate (OCR) was measured using a Seahorse XF-96 analyzer. Before OCR measurement, the sensor cartridge was calibrated with calibration buffer at 37°C in a non-CO_2_ incubator overnight. A 72-h culture was used unless indicated otherwise. For drug treatments, drugs were added 12 h before analysis. Cultures were isolated, trypsinized for 5 min using TRYPLExpress, counted and 200,000 Cells were seeded onto the 96 well microplate coated with Poly-L-Lysine, washed with XF assay medium (pH 7.4) supplemented with 1 mM pyruvate, 2 mM glutamine, and 10 mM glucose and placed in a 37°C incubator without CO_2_ for 1 h before calibration. 1 μM oligomycin, 2 μM protonophoric uncoupler (FCCP) and 0.5 μM antimycin A + rotenone were preloaded in reagent Ports-A, B and C. Three readings were taken after the addition of each reagent prior to injection with subsequent reagents. OCR and ECAR under the same conditions were automatically calculated and recorded by the sensor cartridge and Seahorse XF-96 software. The numerical data of samples in the test were normalized to the protein amount in each well.

### Protein isolation and Western blot analysis

Cells were lysed for 15 min on ice with cell lysis buffer containing 10 mM Tris, 1 mM EDTA, 400 mM NaCl, 0.1% NP-40, 10 µl/ml of phosphatase inhibitors 2 and 3 and 10 µl/ml of protease inhibitor cocktail (MilliporeSigma #P8340). Samples were centrifuged at 12,700 rpm for 15 min at 4℃. Protein was quantified by Bradford assay kit (Bio-Rad #5000001), and 20 µg was separated on SDS-PAGE and transferred to Nitrocellulose membranes. After blocking with 5% non-fat dry milk solution for 1 h, the membrane was incubated with the indicated antibodies overnight. Following three washes with TBST (TBS containing 0.05% Tween-20), appropriate secondary antibody was added for 1 h followed by three washes before scanning with the iBRIGHT 1500 Imaging System (Thermo Fischer Scientific). Antibodies were obtained from the following sources: Cell Signaling Technology (ID1, #23369S; ID3, #9837: RRID AB_2732885); DHSB (A-Actin, # 12G10).

### Xenograft tumors and experimental metastasis assays

To generate primary tumors, SUM149-GFP-Luc cells(7) (2 x 10^6^) were orthotopically injected into NSG mice (RRID: BCBC_4611) (8-12 weeks old) and tumors at end point of about 2000 mm^3^ tumor volume were collected and frozen for proteomics. For electron microscopy and array tomography, a small tumor of about 60 mm^3^ was collected and fixed immediately (for details see sample preparation for array tomography). For experimental lung metastases, mice were randomized to groups by weight and about 0.5 million cells in 100 μl PBS were injected by tail vein into 8-12-week-old NSG female mice. Within two hours, bioluminescence was determined to assure successful inoculation (0-day measurement). Bioluminescence was monitored at 2-week intervals for up to 20 weeks using the IVIS Spectrum imager (PerkinElmer Inc.).

All experiments were according to approved protocols. Animal care was provided in accordance with the procedures outlined in the “Guide for Care and Use of Laboratory Animals” including those pertaining to studies of neoplasia (National Research Council, 1996; National Academy Press; Washington, D.C.). NCI-Frederick is accredited by AALACi and follows the Public Health Service Policy for the Care and Use of Laboratory Animals.

### Sample preparation for proteomic analysis

#### Tissue/Cell lysis, digestion, TMTpro labeling

Three-day cultures from 2D, EmC and SphC from SUM149 and IBC-3 cells were harvested and washed with PBS (2X) and the pellets and/or tissue samples were treated with 300 µl of EasyPep lysis buffer (ThermoFischer Scientific #A45735) and 1 µl of universal nuclease (Thermo #88700). Homogenized with pestle and centrifuged for 5 min at 14,000g. Removed the supernatant to a clean tube. Cell pellets were treated with 100 µl of EasyPep lysis buffer and 1 µl of universal nuclease. Protein concentration was determined using the BCA method. For each sample, the volume corresponding to 50 µg of protein was adjusted to 100 µl total with lysis buffer then treated with 50 µl of reducing and alkylating solutions provided with the EasyPep kit (Thermo #A40006) and incubated at 95°C for 10 min then allowed to cool to room temperature. Samples were then treated with 4 µg of trypsin/LysC provided with the EasyPep kit and incubated at 37°C for 19 h overnight. Samples were then treated with 40 µl of 12.5µg/µl TMTpro in acetonitrile and incubated at room temperature for 1h. Samples were quenched with 50µL of 5% hydroxylamine, 20% formic acid (FA) for 10 min and then combined. Samples were cleaned using spin column provided with the EasyPep kit and eluted peptides were dried in a speedvac.

##### LC-MS analysis

Dried peptides were resuspended in 0.1%FA and injected in triplicate using a Dionex U3000 RSLC in front of a Orbitrap Eclipse (Thermo) equipped with an EasySpray ion source. Solvent A consisted of 0.1%FA in water and Solvent B consisted of 0.1%FA in 80%ACN. Loading pump consisted of Solvent A and was operated at 7 µl /min for the first 6 min of the run then dropped to 2 µl /min when the valve was switched to bring the trap column (Acclaim™ PepMap™ 100 C18 HPLC Column, 3μm, 75μm I.D., 2cm, PN 164535, Thermo) in-line with the analytical column (EasySpray, 2μm, 75μm I.D., 50cm, PN ES903, Thermo). The gradient pump was operated at a flow rate of 300 nl/min and each run used a linear LC gradient of 5-7%B for 1min, 7-30%B for 134 min, 35-50%B for 35 min, 50-95%B for 4 min, holding at 95%B for 7 min, then re-equilibration of analytical column at 5%B for 17 min. All MS injections employed the TopSpeed method with a 3 sec cycle time that consisted of the following: Spray voltage was 1800V and ion transfer temperature of 275 ⁰C. MS1 scans were acquired in the Orbitrap with resolution of 120,000, AGC of 4e5 ions, and max injection time of 50ms, mass range of 400-1600 m/z; MS2 scans were acquired in the Ion Trap using turbo scan method with AGC of 1e4, max injection time of 35ms, CID energy of 35%, isolation width of 0.7Da, intensity threshold of 1e4 and charges 2-5 for MS2 selection. Real Time Search (RTS) was enabled for TMTpro labeled peptides using the Uniprot Human Feb 2020 database. RTS settings were Full tryptic peptides, 1 max missed cleavage, fixed carbamidomethyl on Cys, fixed TMTpro on Lys and peptide N-terminus, and variable oxidation on Met. MS3 scans for peptides identified by RTS were acquired in the Orbitrap with resolution of 50,000, AGC of 2.5e5, max injection of 86ms, mass range 100-500 m/z, and isolation width of 2Da. Advanced Peak Determination, Monoisotopic Precursor selection (MIPS), and EASY-IC for internal calibration were enabled and dynamic exclusion was set to a count of 1 for 15sec.

#### Database searching and post-processing

All three injections were batched together as fractions and all MS files were searched with Proteome Discoverer 2.3 (RRID:SCR_014477) using the Sequest node. Data was searched against the Uniprot (RRID:SCR_002380) Human database from Feb 2020 using a full tryptic digest, 2 max missed cleavages, minimum peptide length of 6 amino acids and maximum peptide length of 144 amino acids, an MS1 mass tolerance of 10 ppm, MS2 mass tolerance of 0.6 Da, variable oxidation on methionine (+15.995 Da) and fixed modifications of carbamidomethyl on cysteine (+57.021), TMTpro (+304.207) on lysine and peptide N-terminus. Percolator (RRID:SCR_005040) was used for FDR analysis and TMTpro reporter ions were quantified using the MS3 Reporter Ion Quantifier node and normalized on total peptide intensity of each channel. TMTpro channel assignment for conditions can be found in Table S2. Statistical analysis was performed within PD2.3 and Log2FC was calculated based on the median of each group and adjusted p-value was calculated using ANOVA. Differentially expressed proteins were set using Log2FC cutoff of +/- 0.6 and adjusted p-value of <0.05. For proteins that were observed in 2 or more replicates of a condition and 0 of the other in a comparison, the Log2FC was set to +/- 0.1 the highest/lowest value in the comparison and adjusted p-value set slightly lower than the lowest value calculated in the comparison. Pathway enrichment analysis was performed using Ingenuity Pathway Analysis (RRID:SCR_008653) and the Log2FC threshold was lowered to +/- 0.1 and the adjusted p-value was kept at <0.05. Data have been deposited at MassIVE (https://massive.ucsd.edu/; RRID:SCR_013665) with accession number MSV000099222.

### Reversed-Phase Ion-Pairing LC-MS^2^ Assay for Measuring Cell Central Carbon Metabolites

Three-day cultures of SUM149 and IBC-3 cell under 2D, SusC, EmC, and SphC conditions were used. All targeted central carbon metabolite (CCM) reference compounds, including acetyl-CoA and α-ketoglutarate, were purchased from Sigma-Aldrich (St. Louis, MO). The stable isotope-labeled internal standards (SI-CCM) were ^13^C_3_-lactate and ^13^C_4_-succinic acid (Cambridge Isotope Laboratory, Andover, MA), as well as ^13^C_6_-glucose-6-phosphate and ^13^C_6_-fructose-1,6-diphosphate (Medical Isotopes, Inc., Pelham, NH). All CCM and SI-CCM analytical standards have reported chemical and isotopic purity ≥ 98% and were used without further purification. OmniSolv^®^ LC-MS grade acetonitrile and methanol were obtained from EMD Millipore (Billerica, MA). Tributylamine (TBA), LC-MS grade acetic acid and formic acid were purchased from Fisher Scientific (Hampton, NH). LC-MS grade ammonium formate was obtained from CovaChem (Loves Park, IL). All chemicals and solvents used in this study were HPLC or reagent grade unless otherwise noted. The isotope dilution liquid chromatography-tandem mass spectrometry method adapted from previous publications (22,23) was employed to measure the concentrations of cell CCMs.In brief, cell pellets were extracted with chilled 80% methanol in water supplemented with appropriate isotopic standards. Reversed-phase ion-pairing LC-MS^2^ analysis was performed using Thermo TSQ Quantiva triple quadrupole mass spectrometer (Thermo Scientific, San Jose, CA) coupled with Shimadzu 20AC-XR liquid chromatography system for the CCM measurement. The mass spectrometers were operated in negative ion mode and set to monitor parent-product ion transitions using Selected Reaction Monitoring (SRM). Quantitation of targeted metabolites was performed using Xcalibur™ Quan Browser (Thermo Scientific). Calibration curves for each metabolite were constructed by plotting CCM/SI-CCM peak area ratios obtained from the calibration samples versus their CCM concentrations and fitting the data using linear regression with 1/*X* weighting. The analyte concentrations in experimental samples were then interpolated using the linear function obtained from the calibration curve.

### RNA isolation and quantitative RT-PCR

Total RNA was isolated using GeneJET RNA purification kit (Thermo Scientific, #K0732), and 2 µg RNA was taken for cDNA synthesis using Superscript Reverse Transcriptase III (RT) according to manufacturer’s instructions (Invitrogen, #18080044). Quantitative PCR was carried out with Fast SYBR Green master mix (#4385612, Applied Biosystems, Foster City, USA) using the 7500 Fast Real**-**Time PCR instrument (Applied Biosystems) and the relative expression levels were measured using the relative quantitation ΔΔ*C*t method and normalized to *actin*. Data represent three independent biological replicates; each measured as technical triplicates. Primers were as follows: *ID1* (5’-aaacgtgctgctctacgaca-3’ and 5’-gattccgagttcagctccaa-3’; *MRPS12 (*5’-gctacctgctccatggctac-3’ and 5’-cacttgcgattggctgagt-3’); *NDUSF6 (*5’-cacactggccaggtttatga-3’ and 5’-tggtgctgtctgaactggag-3’); *RPLP0* (5’-gcaatgttgccagtgtctgtc-3’ and 5’-gccttgaccttttcagcaagt-3’).

### Single Cell Library Preparation, Sequencing and Analysis Method

Three-day cultures of SUM149 and IBC-3 cells grown under 2D, EmC, and SphC conditions were harvested, washed with PBS, and had RNA isolated as per manufacturer’s instructions and 2 µg RNA was taken. Two biological replicates were labeled with Hashtag1 or Hashtag2 for each experimental condition. The single-cell gene expression libraries were generated employing the 10x Chromium Single Cell 3’ Reagent kit v3 (CG000183 Rev A) with Feature Barcode technology in accordance with the guidelines provided by 10X Genomics, US. The TotalSeq -B anti-human Hashtag reagents (BioLegend #394602, 394604, 394606) were used for cell hashing and multiplex samples. These libraries were sequenced on a NovaSeq system utilizing two SP flowcells from Illumina, US. The sequencing run was setup as 28 cycles for Read1, 8 cycles for i7 index, and 75 cycles for Read2 according to protocol recommendation. Sequencing run demultiplexing was performed by using Cellranger mkfastq, with one mismatch allowed in the barcodes. Cell Ranger v4.0.0 was applied for alignment, tagging, gene and transcript counting for Chromium 3’ gene expression libraries, and sample demultiplexing was done using Cell Ranger v4.0.0 based on hashtag feature barcodes. The human genome GRCH38 reference with Gencode v32 /Ensembl98 annotation (refdata-gex-GRCh38-2020-A; RRID:SCR_014966) were used as reference in Cell Ranger analysis. Seurat (v5) (24) R package was used for single cell RNA-seq (scRNA-seq) data analyses, including cell hashtag filtering, quality control, normalization, clustering, visualization, as well as differential expression (DE) analysis. After cell hashtag demultiplexing, only singlets with reasonable expression matrices were kept for downstream analysis. For DE analysis, we conducted pairwise comparisons among the three culture conditions, and we also compared each culture condition versus the other two conditions. Differentially expressed genes (DEGs) were identified with adjusted P value less than 0.05 and average log2 fold change more than 1 (or less than -1) cutoffs. Genes commonly up- or downregulated in both cell lines were used as inputs for QIAGEN Ingenuity Pathway Analysis (QIAGEN Inc., https://digitalinsights.qiagen.com/IPA, version 01-23-01; RRID:SCR_008653) (25). Data are available at GEO under accession number GSE30927.

### Sample preparation for electron microscopy

SUM149 breast cancer cells were cultured as described above, harvested and washed once with PBS, and Karnovsky’s fixative (4% formaldehyde and 2% glutaraldehyde in 0.1 M cacodylate buffer) was added and incubated at RT for 5 h with gentle pipetting every hour to prevent clumping. Samples were centrifuged at 500 rpm for 30 sec and the supernatant decanted. 1 ml cacodylate buffer (0.1 M) was added and centrifuged again at 500 rpm for 30 sec. The pellet was suspended in cacodylate buffer and kept at 4°C. Cells from 2D culture were pelleted and embedded in 1% low-melt agarose before continuing with sample preparation. All other cell growth types were left free-floating. Cells were post-fixed in 2% osmium tetroxide and 1.5% potassium ferricyanide in 0.1 M sodium cacodylate for 1 h at room temperature, washed with ultrapure water, and then stained with 1% aqueous uranyl acetate overnight at 4°C. After further washes in ddH_2_O, the cells were treated with lead aspartate at 65°C for 30 min and washed again. The samples were dehydrated through graded ethanol (35%, 50%, 70%, 95%, and 100% x 5, 10 min. each) and propylene oxide (PO; 5 x 10 min.). The cells were infiltrated with increasing concentrations of Polybed 812 resin (hard formulation) in PO (resin: PO; 1:3, 1:2, 3:1, and 100%), embedded in 100% degassed resin, and cured at 65°C for 48 h in flat BEEM capsules (EMS, Warrington, PA, USA). After polymerization, resin blocks were trimmed with a Leica EM TRIM2 milling system. After trimming the block face down, 100 nm sections were cut with a 45° ultra-diamond knife (Diatome) using a Leica ARTOS ultramicrotome. Sections were collected with a loop and placed on glow-discharged ITO coverslips.

### Array tomography SEM imaging and segmentation

Sections on ITO coverslips were mounted onto 4-inch type-p silicon wafers using copper tape and affixed to a 4-inch stage-decel holder in the GeminiSEM 450 (Carl Zeiss). Low-resolution overview scans of individual resin sections were acquired at 150 nm pixel resolution using ATLAS 5 Array Tomography software (Fibics). High-resolution (5 nm pixels) large areas were imaged by stitching multiple fields, enabled by automated stage movement and a four-quadrant backscatter detector (aBSD, Zeiss). Image strips of roughly 40,000 x 8,000 pixels were cropped from the larger images to capture roughly similar numbers of cells. These strips were imported into Napari (26) where segmentation was performed using the empanada plugin. The appropriate deep learning models were deployed in the 2D inference module with image tiling: mitochondria were segmented using MitoNet_v1 (27) lipid droplets with DropNet_base_v1, and nuclei with NucleoNet_base_v1 (to be reported elsewhere). Errors in model predications were corrected manually in empanada-napari. To facilitate correction in these large images, the “create tiles” module in empanada was used to generate smaller image tiles to reduce the time needed for manual adjustments. After all corrections were completed on the individual tiles, both the images and their segmentations were reassembled using the “merge tiles” module in empanada. Cell segmentation was performed in 3DSlicer (Ver. 5.8.1; (28) using thresholding to delineate borders for each cell in every image, followed by filling in using the fill tool in Napari. Quantitative measurements of cells, mitochondria, lipid droplets, and nuclei were obtained with the napari-clusters-plotter plugin, which utilizes region props from scikit-image (29, 30). The cell packing ratios were determined by taking the total pixel area of all cells that were segmented in an image divided by the total pixel area of the field of view.

### Statistics

Unless stated otherwise, quantitative data were analyzed by the two-tailed unpaired t test and are shown as the mean ± S.E.M. Graphs were made using GraphPad Prism10 (RRID:SCR_002798) software. The number of samples (n) refers to biological replicates unless indicated otherwise.

## RESULTS

### Durable alterations of breast cancer cell proliferation in 3D depending on culture paradigm

In this study, we examined four culture conditions (**Figure 1A)** using three epithelial breast cancer cell lines (SUM149, IBC-3, and MDA-MB-468), which were chosen after comparing several breast cancer cell lines for their ability to form 3-dimensional assemblies in EmC (Figure S1A). In contrast to the original report of this culture method (19), we observed that all cell lines formed “emboli”, with those by basal epithelial cell lines appearing most compact. Apart from conventional 2D culture of these adherent cells on plastic, all other conditions were in ultra-low attachment plates. Because emboli culture (EmC) requires addition of PEG8000 to standard, cell-line specific media (19), PEG8000 was also added to 2D and suspension culture (SusC) to control for the effects of media viscosity and molecular crowding. Thus, comparison of 2D and SusC reveals the effect of losing substrate attachment. Comparison of SusC and EmC provides information on the effect of the dynamic nature of EmC, which is the only difference between these two types of culture. The media composition is the only difference between SusC and Sphere cultures (SphC) which were in specific medium that enriches stem cell like features (31). EmC and SphC differ in both media composition and dynamic versus static conditions. Bright-field imaging of 3-day cultures revealed characteristic structures formed under each 3D condition (**Figures 1B and S1A**). Cell clusters were smallest in SusC, where loss of substrate attachment limits aggregation, largest in EmC, likely reflecting enhanced aggregation and reinforced cell-cell adhesions under fluid dynamic forces. Comparison of the numbers of cells within 3D structures (isolated by brief low-speed centrifugation suggested that loss of substrate by SusC reduced the cell proliferation rate, whereas the media in SphC rescued proliferation, likely because it is optimized for stemness (**Figure 1C-D**). Analysis of cell cycle stages by DNA content in SUM149 and IBC-3 cells supported this conclusion and indicated that cell death (sub-G1 DNA content) was minimal. The mechanical forces in EmC did not change the proliferation rate of SUM149 or MDA-MB-468 cells in suspension, but reduced proliferation of IBC-3 cells even further. Notably, when cells from 3D culture were dissociated and re-plated in 2D, these proliferation rate differences persisted for at least 4 days (**Figure 1E**). Compared to cells never exposed to 3D culture, cells from SphC proliferated more rapidly, whereas cells from SusC, and especially EmC, showed a durable reduction in their proliferation rates. These results indicate that 3D culture conditions impart lasting cell adaptations that are detectable at the level of proliferation. To address molecular differences, we conducted single-cell mRNA-Seq (scSeq) of SUM149 and IBC-3 cells in 2D, SphC and EmC. UMAP cluster analysis revealed that each population of SUM149 cells segregated well with only limited overlap of clusters containing cells of two or more culture conditions (**Figure S2A-B**). For IBC-3, cells in 2D and EmC were more similar, while those in SphC segregated the most. Next, we identified differentially expressed genes (DEGs) for each condition in bulk to all others (**Supplemental File 1**) and performed pathway analyses (**Supplemental File 2**). Comparison of the pathways with most significant z-scores validated the observed reduction in cell proliferation in EmC while cholesterol biosynthesis was upregulated in SphC (**Figure 1F**). Increased cholesterol biosynthesis may be consistent with the higher proliferation rate in SphC, although mitosis-related pathways were only upregulated in IBC-3 in SphC (Supplemental File 2). Collectively, these results demonstrate that cancer cells durably and differentially alter their cell proliferation rate in SphC versus EmC.

**Figure 1.**
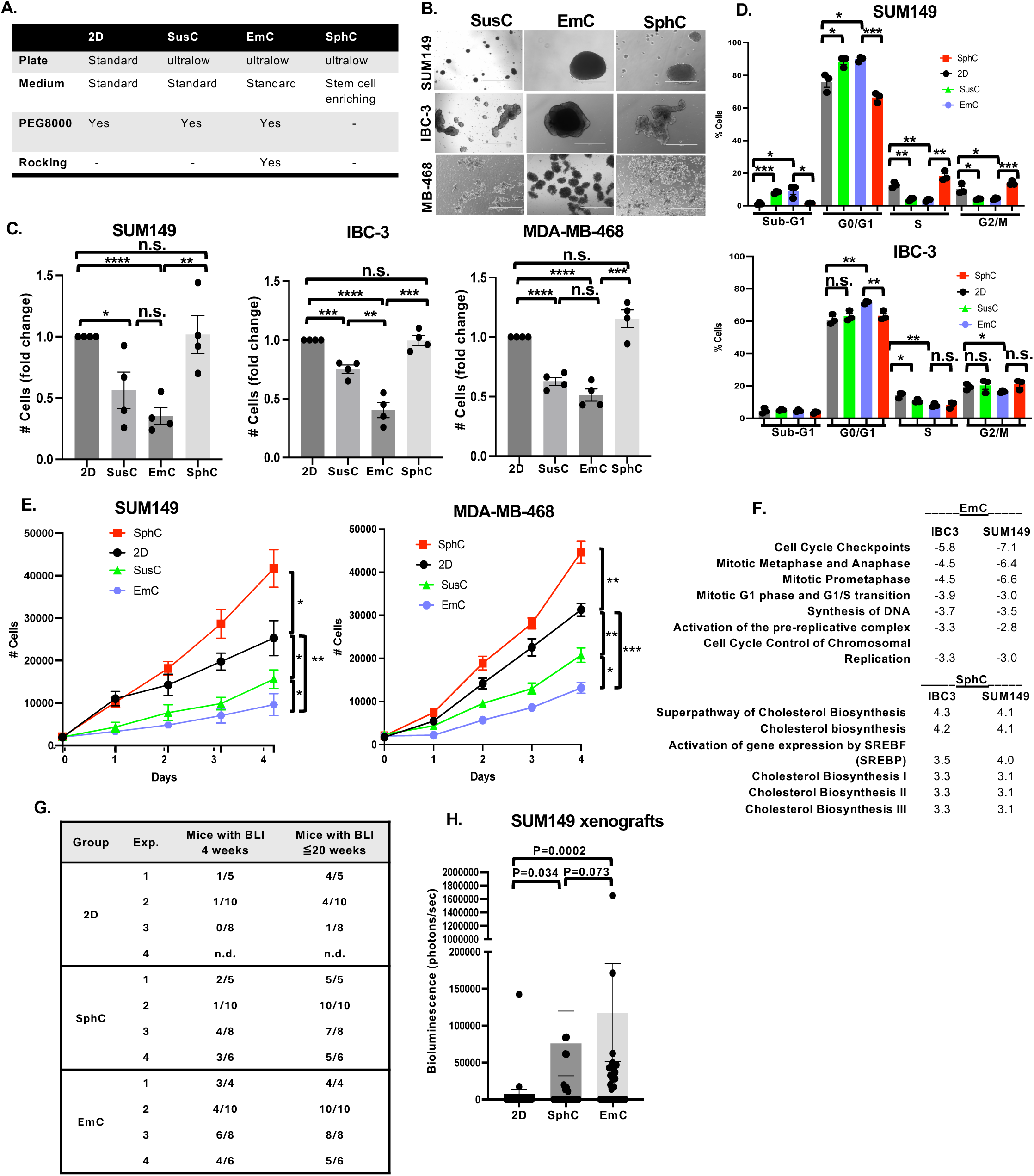
Cells cultured in EmC reduce cell proliferation ex vivo but acquire lung colonization efficiency in vivo. **A,** Overview of cell culture conditions used in this study (see Methods for further details). **B,** Brightfield images of the of indicated cell lines after three days in the specified 3D culture conditions (scale bar =1 mm). **C,** Viable cell counts after three days of culture under the indicated conditions, all starting with equal cell numbers (n = 4). **D,** Flow cytometric analysis of DNA content of the cells from SUM149 and IBC-3 cells cultured as in panel C (n = 3). **E,** Quantification of viable cells by imaging cytometry replated in 2D (day 0) after 3 days under the indicated culture conditions (n = 3). Statistical analysis refers to day 4. **F,** Z-scores with P<0.05 derived from IPA analysis of common DEGs in EmC and SphC respectively compared to all other conditions. **G,** Number of mice across four independent experiments, showing lung bioluminescence (BLI) after tail vein injection of SUM149 cells that had been cultured as 2D, SphC and EmC for 3 days before injection by tail vein. Data are for measurements at week 4 and cumulative numbers until week 20 after injection. *P* values determined by Fischer’s exact test as follows:. 2D vs SphC, *P*= 0.045; 2D vs EmC, *P*= 0.00013; SphC vs EmC, *P*= 0.065. **H,** Quantification of bioluminescence of all mice at week 4 post-injection (n = 22-29). *P* values were determined by Wilcoxon Test (unpaired, two-sided). All quantitative data, except panels G-H, are mean ± SEM, \**P*<0.05, \*\**P*<0.01, \*\*\**P*<0.001, \*\*\*\**P*<0.0001; n.s., not significant.

Mammosphere culture is known to enrich for cancer stem cells in cell lines, increasing the tumor take rate and experimental metastasis rate (32,33). In contrast, the reduced proliferation rate of cells in EmC could potentially indicate senescence or dormancy. Pilot experiments determining these states were inconclusive, so we next assessed the capacity of SUM149 cells, which express a luciferase reporter, to seed experimental metastasis. Cells from 3D cultures were dissociated into single cells and injected by tail vein into immunocompromised mice; 2D-cultured cells served as controls. Bioluminescence intensity (BLI) was determined on the same day to confirm lung seeding. As anticipated, most injected cells died during the initial phase post-injection, and four weeks later most mice did not show any BLI (**Figure 1G**). However, as expected (33), the frequency of mice with BLI signal was higher in mice which received cells from SphC (34.5%) compared to 2D controls (8.7%). In pilot experiments, we had determined that injection of 10^5^ cells was suitable to demonstrate the enhanced colonization by cells from SphC. Strikingly, even more mice (60.7%) injected with cells from EmC exhibited lung colonization, and BLI quantification confirmed that EmC increased lung colonization efficiency relative to both 2D or SphC cells (**Figure 1H**). By week 20, nearly all mice injected with EmC-derived cells developed detectable lung colonies (**Figure 1G**). While we cannot exclude the possibility that re-establishment of cell-cell adhesions between cells from EmC cultures enhanced initial survival after injection (33), these results demonstrate that EmC does not diminish malignancy. Rather, despite their lower proliferation rate in culture, EmC-derived cells exhibit heightened metastatic competence.

### SUM149 ultrastructural features in EmC bear most resemblance to primary tumors

Given the distinct morphologies of cell assemblies across the three 3D conditions (Figure 1B), we employed volume electron microscopy, specifically array tomography (34), to assess the ultrastructural adaptations of SUM149 cells and their resemblance to *in vivo* tumor tissue. As shown in **Figure 2A**, electron microscopy images revealed that the cell packing ratio was highest and most similar in EmC and tumor tissue. Cells from 2D culture were included in this analysis, but packing ratio does not apply to this condition. In addition, mitochondria and nuclei appeared largest, and cell-to-cell connectedness most uninterrupted, in EmC and tumors (**Figures S3A-B**). AI-based algorithmic quantification of nuclear and mitochondrial morphometrics in one sample per condition further underscored these similarities. Nuclear analyses showed that area and solidity, signifying mostly convex or positive curvature of the membrane, were highest and most similar in tumor and EmC samples (**Figure 2B**). Nuclear aspect ratios (length/width) and perimeters were similar across culture conditions and tumor tissue (**Figure S3C**). The data indicate that nuclear morphology in EmC resembles the more “normalized” phenotype of tumors, though increased nuclear size has also been correlated to neoplastic malignancy (35). Mitochondrial analysis revealed that cells in EmC and tumor tissue harbored the largest mitochondria by area and major axis length, whereas mitochondria in SphC were the least circular with the highest aspect ratios (**Figures 2C and S3D**). To extend this analysis, we developed new algorithms to examine lipid droplet (LD) morphology. Cells in SphC contained the largest LDs with the greatest area and major axis length, while LDs in EmC more closely resembled those in tumor tissue, showing similarity in major axis length, circularity, and staining intensity (**Figures 2D and S3E-F**). Although the determinants of LD staining intensity remain unclear, we speculate that the lipid composition could be a determining factor. In contrast to a prior ultrastructural study of SUM149 emboli (36), we did not observe many microvilli. Full quantifications and statistical analyses are provided in **Supplemental Files 3 and 4**. Together, these data suggest that SUM149 cells adopt distinct ultrastructural features depending on culture condition, and that EmC recapitulates certain aspects of mitochondrial, nuclear, and lipid droplet morphology observed in tumor tissue *in vivo*.

**Figure 2.**
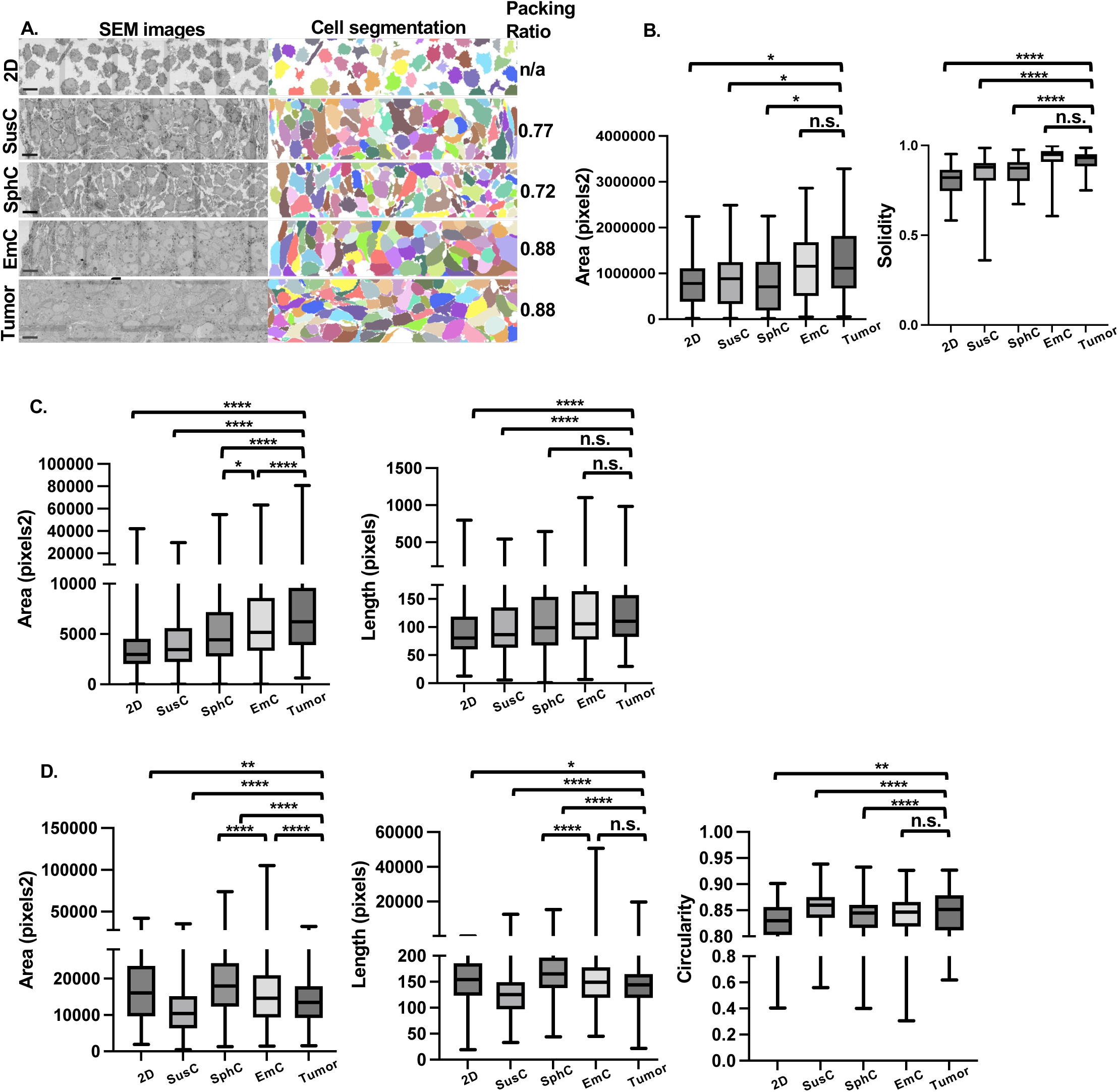
Ultrastructural analysis of SUM149 cell cultures and tumor tissue. **A,** Representative scanning electron microscope images and corresponding cell segmentation of SUM149 cells after 3 days in the indicated culture conditions, as well as from a xenograft tumor. Scale bar = 10 µm. **B,** Nuclear metrics of the indicated features for each condition (n = 50-68). **C,** Mitochondrial metrics of the indicated features for each condition (n = 1451-1995). **D,** Lipid droplet metrics of the indicated features for each condition (n = 131-654). \**P*<0.01, \*\*\*\**P*<0.00001; n.s., not significant.

### Adaptation of mitochondrial metabolism depends on 3D paradigm

To probe molecular adaptations to different growth conditions, we conducted proteomic analysis of the total levels of proteins in 2D, SphC, and EmC compared to xenograft tumor tissues (**Supplemental File 5**). Principal component analysis (PCA) plots of the data from culture conditions (2D, SphC, and EmC) and primary tumors showed well separated clustering of the biological replicates (**Figure S4A**). Using the 2D proteome as reference, we compared the differentially expressed proteins (DEPs) between conditions (**Figures 3A and S4B**). As expected, the more complex and heterogenous tumor tissue was most divergent from cells cultured in 2D (236 up, 576 down). Cells from SphC showed the least DEPs (70 up, 15 down), while cells from EmC were intermediate (148 up, 159 down). Cells from EmC showed the most overlap with tumor tissue (**Figure 3A)**. Analysis of the proteins exclusively overrepresented in EmC and tumors by ToppGene (37) showed that 35/39 were part of the Metabolism Reactome (GSEA ID MM14563), while 116/132 underrepresented proteins were part of the RNA Metabolism Reactome (GSEA ID M16843), encompassing splicing and translation. Ingenuity Pathway Analysis (**Figures S4C and Supplemental File 5**) further showed that translational activity was underrepresented in tumors and in EmC consistent with the reduced proliferation rate in EmC compared to 2D (Figure 1D). Regarding metabolic pathways, not surprisingly, tumor contained more proteins in glycolysis and less in oxidative phosphorylation pathways (**Figure 3B**). In contrast, both EmC and SphC showed increases in TCA cycle pathway as well as glycolysis and/or Warburg effect. Collectively, the proteomic analysis showed distinct and common adaptations of cells to different 3D culture conditions and suggest that cells in EmC share more similarities with tumor tissue than do cells in SphC.

**Figure 3.**
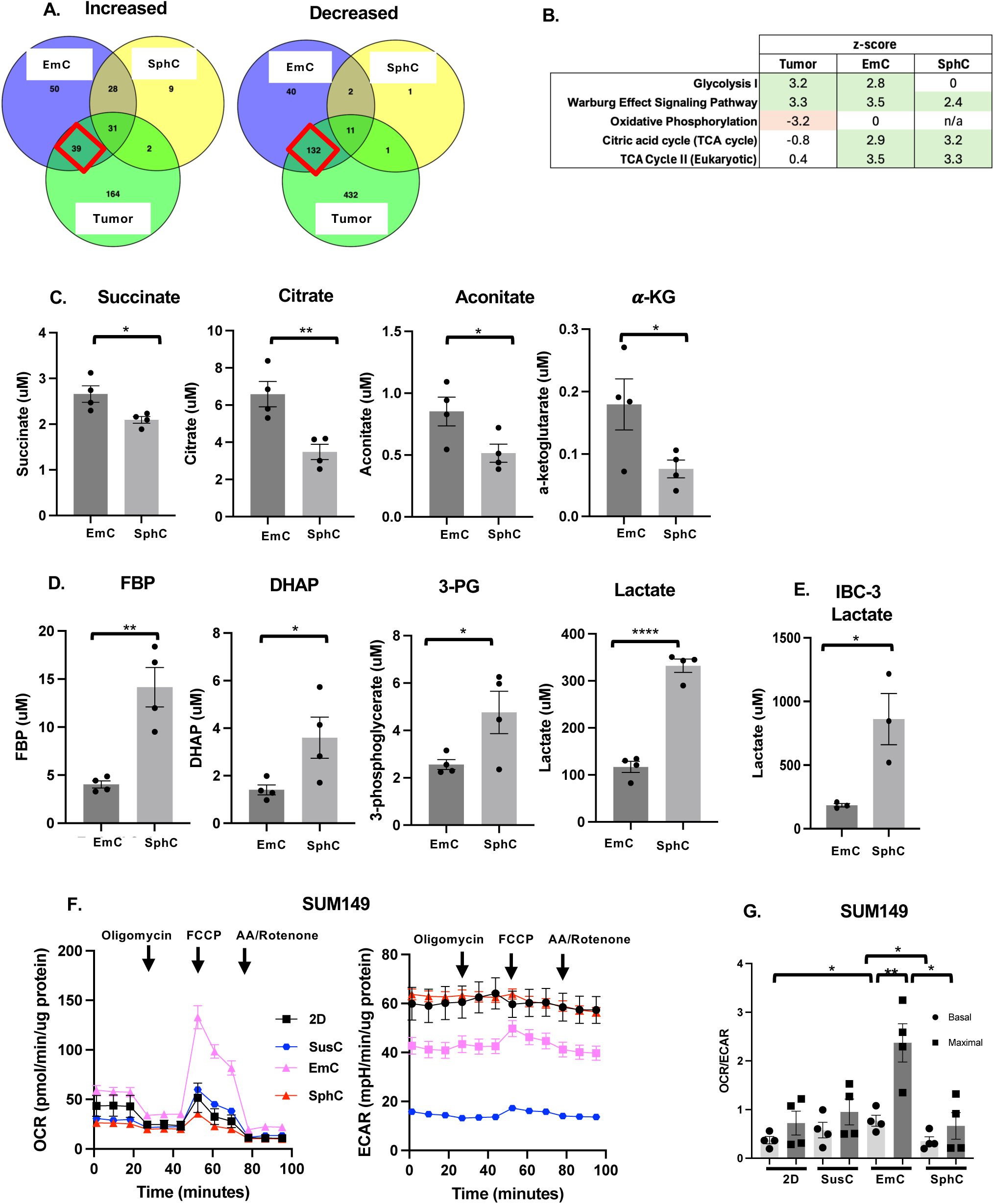
Proteomic and metabolic analysis reveals differential cell adaptations to 3D culture. **A,** Venn diagram showing the numbers of SUM149 proteins differentially under- or overrepresented in EmC, SphC and tumor samples compared to 2D cultures. **B,** Most significant pathway z-scores with P<0.05 in common between primary tumors, EmC, and SphC in comparison to 2D. **C,** Fractional enrichment of the indicated intracellular metabolites of SUM149 cells in SphC and EmC (n = 4). **D,** Fractional enrichment of the indicated intracellular metabolites of SUM149 cells in SphC and EmC (n = 4). **E,** Intracellular lactate levels in IBC-3 cells under the indicated culture conditions (n = 3). **F,** Oxygen consumption rate (OCR) and extracellular acidification rate (ECAR) over time for SUM149 cells derived from the indicated culture conditions. Shown are a representative data normalized to protein, with 3 technical replicates per time point. **G,** OCR/ECAR ratios of SUM149 cells as in panel F, data are from 4 independent experiments each with 3 time points and 3-4 technical replicates each. Bar graphs show means ± SEM, \**P*<0.05, \*\**P*<0.01, \*\*\*\**P*<0.0001; n.s., not significant.

To further examine metabolic aspects, we determined the levels of central carbon metabolites in SUM149 cells by mass spectrometry. Four TCA cycle metabolites were enriched in EmC compared to SphC (**Figure 3C**). Similar observations were made at the level of pyruvate and glutamate, which feeds into the TCA cycle, and NADH, which is largely produced by the TCA cycle and fuels the electron transfer chain (ETC) (**Figure S4D**). In contrast, four glycolysis pathway intermediates including lactate were enriched in SphC in comparison to EmC (**Figure 3D**). Other metabolites that were measured were similar between conditions, namely malate, fumarate fructose-6-phosphate, phosphoenolpyruvate, glucose-6-phosphate and glyceraldehyde-3-phosphate. Elevated levels of lactate in SphC over EmC were also observed in IBC-3 cells (**Figure 3E**). While steady state levels of metabolites alone do not allow conclusions on the activity of metabolic enzymes, these data suggested that TCA cycle activity of cells in EmC may drive oxidative phosphorylation (OXPHOS) while cells in SphC may favor glycolysis. To test this hypothesis directly, we proceeded to functionally examine metabolic activities by Seahorse metabolic flux technology. For this, the 3D cell structures were dissociated and single cells seeded for analysis. The basal and maximal energy metabolism of cells in different culture conditions were assessed by analyzing oxygen consumption rates (OCR) and extracellular acidification rate (ECAR) ratios. OCR is a measure of mitochondrial respiration and oxidative phosphorylation, and ECAR is a measure of glycolytic activity. Data in **Figure 3F-G** showed that SUM149 cells in EmC exhibited the highest basal and maximal OCR/ECAR ratios. IBC-3 and MDA-MB-468 cells demonstrated overall higher metabolic activity. Nevertheless, cells in EmC exhibited the highest OCR/ECAR ratio (**Figures S4E-F**). Collectively, these results suggest that EmC supports OXPHOS, while SphC supports glycolytic metabolism.

### Specific 3D models reveal distinct drug vulnerabilities

Given the differential metabolic pathway activities between cells in EmC versus SphC, we next examined their responses to pathway-specific inhibitors. For OXPHOS, we used IACS-010759, a complex I inhibitor of the ETC (38). In addition, we used sodium dichloroacetate (DCA) which inhibits pyruvate dehydrogenase kinases (PDK) and thereby promotes the conversion of pyruvate to acetyl CoA, blocking its conversion to lactate (39). Treatment of established 3D cultures of SUM149 or IBC-3 cells with IACS-010759 for three days lead to a dose-dependent reduction in cell numbers in EmC (**Figure 4A**), accompanied by increased cell death (**Figure S5A**), whereas SphC cells were completely resistant. Given that IBC-3 cells showed high levels of basal OCR in all culture conditions, the resistance to IACS-010759 in SphC was surprising. MDA-MB-468 cells were not resistant to IACS-010759 in SphC but still less sensitive compared to EmC (Figure 4A). In contrast, cells in SphC showed significant sensitivity to DCA while cells in EmC were comparatively more resistant (**Figures 4B** and **S5B)**. To specifically probe the role of anaerobic glycolysis (Warburg effect) we used the LDH inhibitor FX-11, which was also more effective in SphC compared to EmC in all three cell lines (**Figure 4C**). Taken together, these data indicate that cells in SphC rely mostly on glycolysis, which is consistent with previous reports (39,40). In contrast, cells in EmC are more dependent on OXPHOS for their metabolic sustenance.

**Figure 4.**
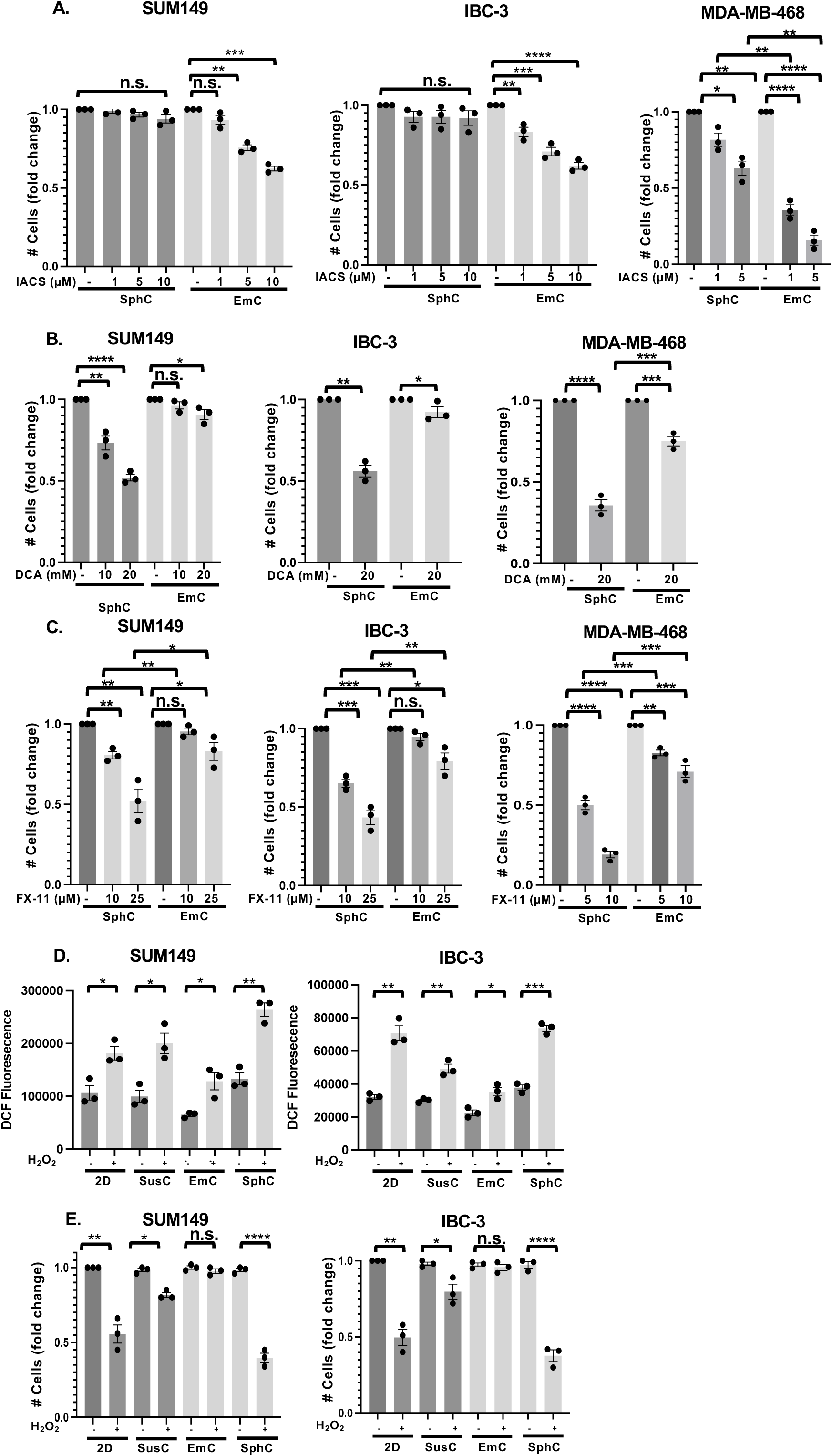
EmC renders cells sensitive to ETC inhibitors and protected from oxidative stress. **A,** Fold change in the numbers of viable SUM149, IBC-3 and MDA-MB-468 cells in EmC and SphC after 72 hours of treatment with OXPHOS inhibitor IACS-010579, added at the indicated concentrations to the established cultures. DMSO was used as vehicle control (n = 3). **B,** Fold change in the numbers of viable SUM149, IBC-3 and MDA-MB-468 cells in EmC and SphC after 72 hours of treatment with DCA, added at the indicated concentrations to the established cultures. DMSO was used as vehicle control (n = 3). **C,** Fold changes in the numbers of viable SUM149, IBC-3 and MDA-MB-468 in EmC and SphC after 72 hours of treatment with FX-11, added at the indicated concentrations to the established cultures. DMSO was used as vehicle control (n = 3). **D,** Quantification of reactive oxygen species (ROS) by DCF fluorescence measurements in SUM149 and IBC-3 cells from the indicated culture conditions. H_2_0_2_ (100 µM) was added to the dissociated cells 15 min prior to analysis (n = 3, mean ± SEM). **E,** Fold change in the numbers of viable SUM149 and IBC-3 cells as in panel C after 2 hours of exposure to H_2_0_2_. Data are expressed as mean ± SEM, \**P*<0.05, \*\**P*<0.01, \*\*\**P*<0.001, \*\*\*\**P*<0.0001); n.s., not significant.

Mitochondrial oxidative phosphorylation is a source of reactive oxygen species (ROS) and may trigger oxidative stress (41). We therefore asked whether cells in EmC may exhibit elevated ROS levels or oxidative stress by staining with the cell-permeable fluorogenic dye H_2_DCFDA (42). Furthermore, we assessed the response to imminent oxidative stress through exposure to exogenous H_2_O_2_. As shown in **Figure 4D**, the basal level of H_2_DCFDA fluorescence of SUM149 and IBC-3 cells was in fact lower in EmC-derived cells in comparison to 2D and SphC. As expected, H_2_O_2_ exposure for 15 minutes increased H_2_DCFDA fluorescence in all culture conditions but remained lowest in EmC. After two hours of exogenous H_2_O_2_ exposure, the numbers of viable cells were reduced in 2D and SphC relative to untreated controls but were unchanged in EmC (**Figure 4E**). These data indicate that, despite their reliance on OXPHOS, EmC cells are most protected from oxidative stress..

### EmC promotes features associated with lung metastasis

Metabolic plasticity is a hallmark of cancer and the metastatic process (43). In epithelial breast cancer, oxidative mitochondrial metabolism and fatty acid oxidation have been associated with metastatic capacity and especially to the lung (44, 45), which is among the most common sites of breast cancer metastasis (46). In EmC, we observed induction of two OXPHOS related genes that are associated with lung metastasis, *NDUFS6* and *MRPS12* (47) (**Figure 5A**). Furthermore, analysis of the scRNA-Seq data showed that other genes associated with lung metastases were also enriched by EmC in SUM149 and/or IBC-3 cells: *ID1, ID3, SPARC, EREG*, *PTGS2* (48–50), *KRT16* (51,52), *MMP7* (53), and *LCN2* (54) (**Figure 5B**). Validation experiments confirmed the upregulation of ID1 at both the mRNA and protein levels in SUM149 and MDA-MB-468 cells (**Figures 5C-D**). ID1, an inhibitor of E-box binding transcription factors, is an important regulator of cell cycle, self-renewal, and tumorigenesis (55–57). Consequently, treatment of cells in EmC with AGX51, which targets ID proteins for degradation (58), reduced ID1 and ID3 protein levels (**Figure 5E**) and decreased viable cell numbers in a dose-dependent manner (**Figures 5F and S5C**). Collectively, these results indicate the relevance for ID1/3 upregulation in EmC and suggest a potential mechanism contributing to the lung colonizing capacity of EmC-derived SUM149 cells (Figure 1G-H).

**Figure 5.**
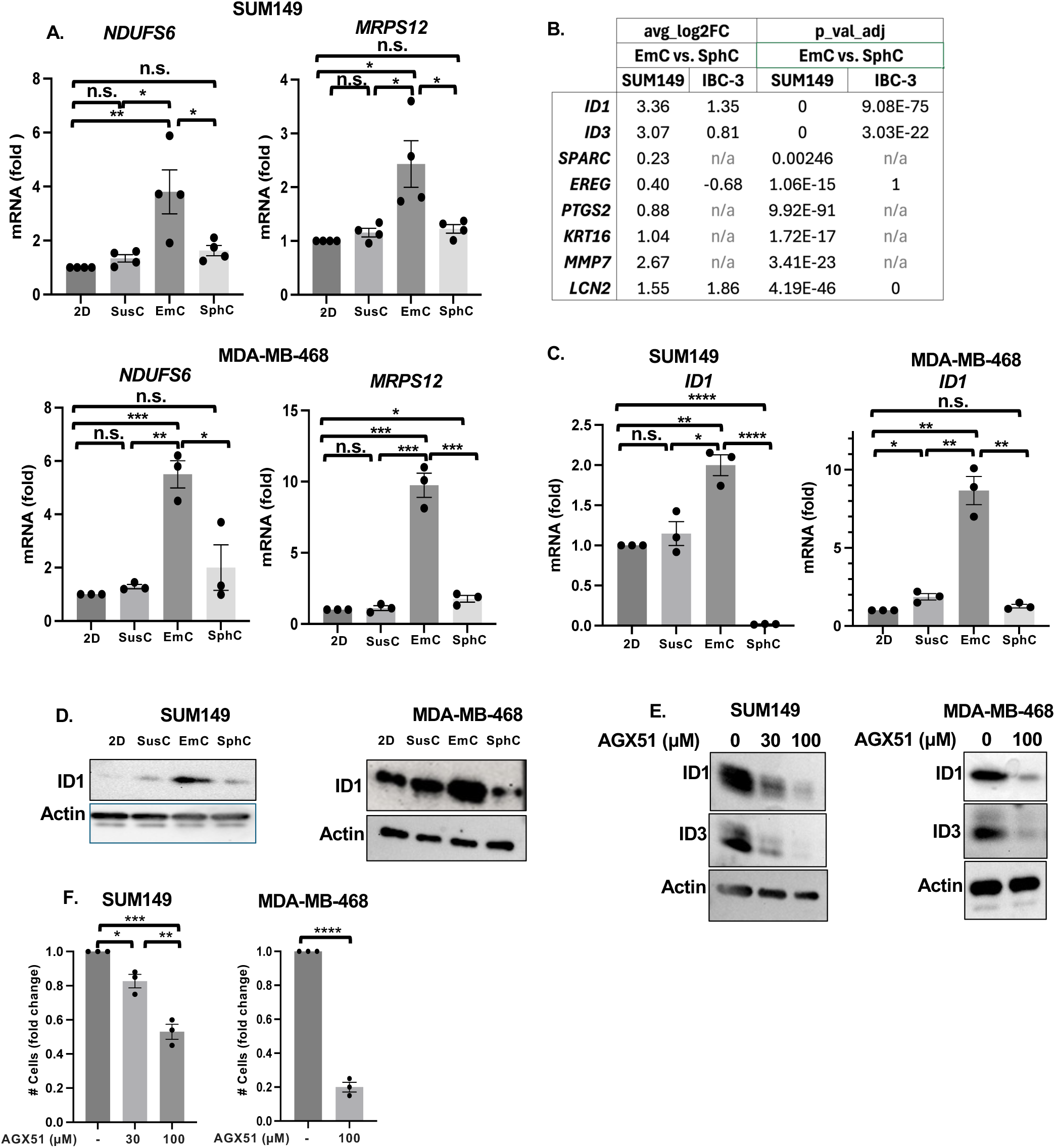
EmC leads to induction of genes related to lung metastasis. **A,** qRT-PCR of mRNA levels for indicated genes in SUM149 and MDA-MB-468 cells under the indicated culture conditions (n = 3). **B,** Differential expression of the indicated genes derived from the scRNA-Seq data comparing EmC versus SphC in SUM149 and IBC-3 cells as indicated. **C,** qRT-PCR (top, n = 3) of *ID1* expression in SUM149 and MDA-MB-468 cultures as indicated. **D,** Western blot analysis of ID1 expression of cells as in C. Actin served as loading control. **E,** Western blot analysis of ID1 and ID3 proteins in established SUM149 EmC that were treated with AGX51 at the indicated concentrations for 3 days. Actin served as loading control. **F,** Fold change in viable SUM149, and MDA-MB-468 cells in established EmC following AGX51 treatment at the indicated concentrations for 3 days (n = 3). Data are shown as mean ± SEM, \**P*<0.05, \*\**P*<0.01, \*\*\**P*<0.001, \*\*\*\**P*<0.0001; n.s., not significant.

## DISCUSSION

The primary objective of this study was to comprehensively characterize breast cancer cells grown in emboli culture (EmC) and compare the findings with more traditional cell culture models such as 2D culture and mammospheres (SphC). Our results indicate three distinct advantages of EmC as an *ex vivo* model system for epithelial breast cancer: (a) reduced cell proliferation, (b) an oxidative mitochondrial metabolic profile (OXPHOS), and (c) ultrastructural and architectural features reminiscent of tumor tissue *in vivo*. These characteristics are crucial because the high proliferation rates in standard *in vitro* systems often make cells artificially sensitive to pharmacological interventions, leading to false positive results in drug screening and ultimately poor clinical translation (4). In contrast, 3D models, including EmC (7), typically exhibit resistance to drug treatments, making them more appropriate for evaluating therapeutic responses (59).

Notably, we observed that the fluid dynamics inherent to the EmC system promote tissue-like cell-cell contacts and ultrastructural changes. While previous research has explored the impact of fluid shear stress and tissue stiffness on cancer progression (60,61), we do not claim that the specific mechanical forces at play in EmC directly model a particular in vivo scenario. However, tissues are exposed to pulsatile fluid flow in blood and lymph vessels, which are likely providing mechanical cues. For lung cancer, specifically, the lung environment provides significant mechanical forces that shape tumor development and progression (62). Consistent with this, our transcriptomic analysis revealed upregulation of several genes in EmC that associated with breast cancer lung metastasis. Among these, keratin 16 (*KRT16*), a regulator of mitochondrial dynamics that is upregulated in lung metastases (51,63), stand out as circulating tumor cells (CTCs) expressing *KRT16* are also associated with shorter relapse-free survival (52). Other lung-metastasis-associated genes were upregulated in EmC, including mitochondrial *MRPS12* and *NDUFS6*, as well as *ID1* and *ID3*, which are implicated in promoting OXPHOS and lung colonization (51). Higher expression of *MRPS12* and *NDUFS6* correlates with decreased metastasis-free survival in breast cancer (47). Interestingly, despite robust mitochondrial oxidative activity, EmC cells maintained low reactive oxygen species (ROS) levels, consistent with previous findings that active TCA cycling is associated with reduced ROS generation (64). This observation aligns with reports that an epithelial phenotype is linked with protection from oxidative stress, potentially mediated through ID1 and KRT16 (63,65).

At the level of energy metabolism, our analyses demonstrate that SphC cells predominately rely on aerobic glycolysis, whereas EmC cells favor mitochondrial OXPHOS. Although glycolytically derived pyruvate canonically feeds the TCA cycle and OXPHOS unless diverted to lactate via the Warburg effect (39,66), recent studies suggest that lactate itself can serve as a fuel for mitochondrial metabolism (67,68). However, only EmC cells, but not SphC cells, were sensitive to electron transfer chain inhibition, underscoring their reliance on mitochondrial oxidative metabolism. These observations are especially relevant in the context of metastasis, where metabolic reprogramming enables tumor cells to adapt and thrive within new microenvironments (69,70). Studies have shown that OXPHOS is induced in epithelial breast cancer cells, specifically CTCs and metastasis-initiating cells (44,71–73). Elevated OXPHOS gene expression is correlated with reduced patient survival, resistance to chemotherapy, and is a metabolic vulnerability in therapy-resistant triple-negative breast cancers (74–77). Consistent with this, metastatic breast cancer cells in the lung amplify OXPHOS metabolism (78), and inhibition of OXPHOS specifically impedes tumor cell seeding of the lung—even when the primary tumor itself is more glycolytic (51).

Combined with our findings that EmC-derived cells have enhanced colonization of mouse lungs and the prevalence of OXPHOS metabolism, the above reports suggest that EmC may serve as a valuable ex vivo paradigm to investigate pathways relevant to lung metastasis. Although most of our analyses were performed using two inflammatory breast cancer (IBC) cell lines, some findings were validated in a non-IBC TNBC line. Additionally, given that our prior work identified a pathway through EmC that was also observed in a non-IBC TNBC PDX model, i.e. COX-2 through AKT/GSK3β to E-cadherin stabilization (7), we postulate that the EmC system may be broadly applicable at least to basal epithelial breast cancers, and potentially other subtypes, although this warrants further investigation. Identifying an *ex vivo* culture model that faithfully recapitulates in vivo tumor phenotypes, especially of difficult to study lung metastases, is critical for dissecting the mechanisms that empower metastatic breast cancer cells and for uncovering new therapeutic targets. Our findings establish EmC as a resource-efficient ex vivo model that captures the ultrastructural, metabolic, and functional hallmarks of metastatic breast cancer, providing a valuable tool for probing the biology and therapeutic sensitivities of other epithelial tumor types.

## Supporting information

Balamurugan et al. EmC 3D Supplementary Figures

## ACKNOWLEDGEMENTS

We are grateful for the superb support provided by the NCI/CCR Flow Cytometry Core and the Laboratory Animal Sciences Program including Small Animal Imaging Program at Leidos Biomedical Research Inc., and Drs. Bao Tran and Yongmei Zhao (FNLCR) for single-cell sequencing. We also thank MaryBeth Hilton and Dr. Brad St. Croix (NCI/CCR) for their kind assistance with NSG mice and Dr. Shiba P. Dash (NCI/CCR) for help with flow cytometry analysis.

This research was supported by the Intramural Research Program of the National Institutes of Health (NIH), ZIA BC 010307, and in part with federal funds under contract no. 75N91019D00024. The contributions of the NIH author(s) were made as part of their official duties as NIH federal employees, are in compliance with agency policy requirements, and are considered Works of the United States Government. However, the findings and conclusions presented in this paper are those of the author(s) and do not necessarily reflect the views of the NIH or the U.S. Department of Health and Human Services.

## Notes

Conflict of interest disclosure statement: The authors declare no conflict of interest

### Competing Interest Statement

The authors have declared no competing interest.

### Summary of Updates

This version includes additional supporting data.

