## Supplementary material for "A simple liquid 3D cell culture paradigm models oxidative mitochondrial metabolism of epithelial breast cancer cells with relevance for lung metastases": Balamurugan et al. EmC 3D Supplementary Figures

Figure S1

A.

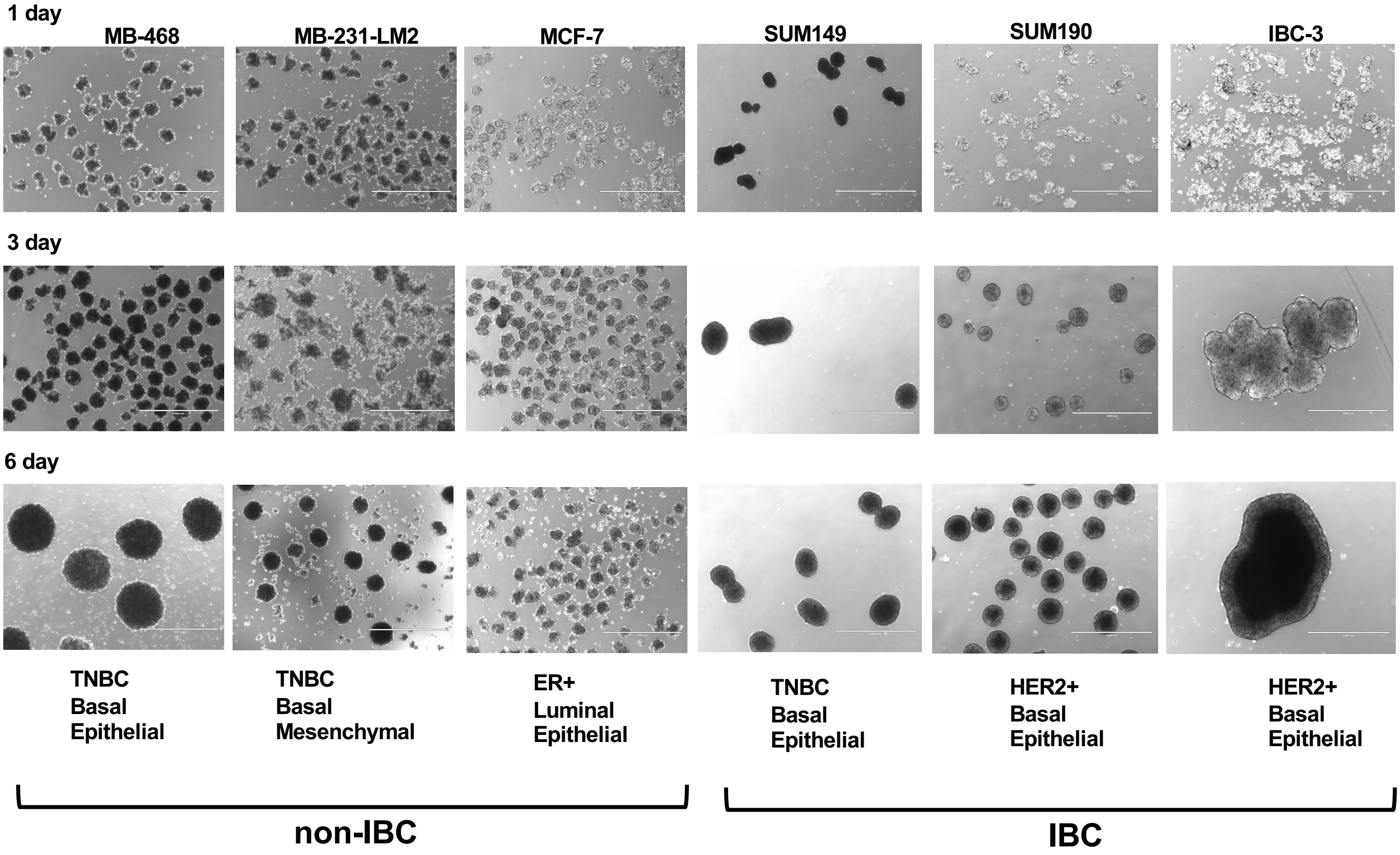


**Figure S1. Emboli formation by different breast cancer cell lines.**

A. Light microscopy images of the indicated IBC and non-IBC cells lines after 1, 3, and 6 days in EmC. Scale bar=1 mm.

**Figure S2**

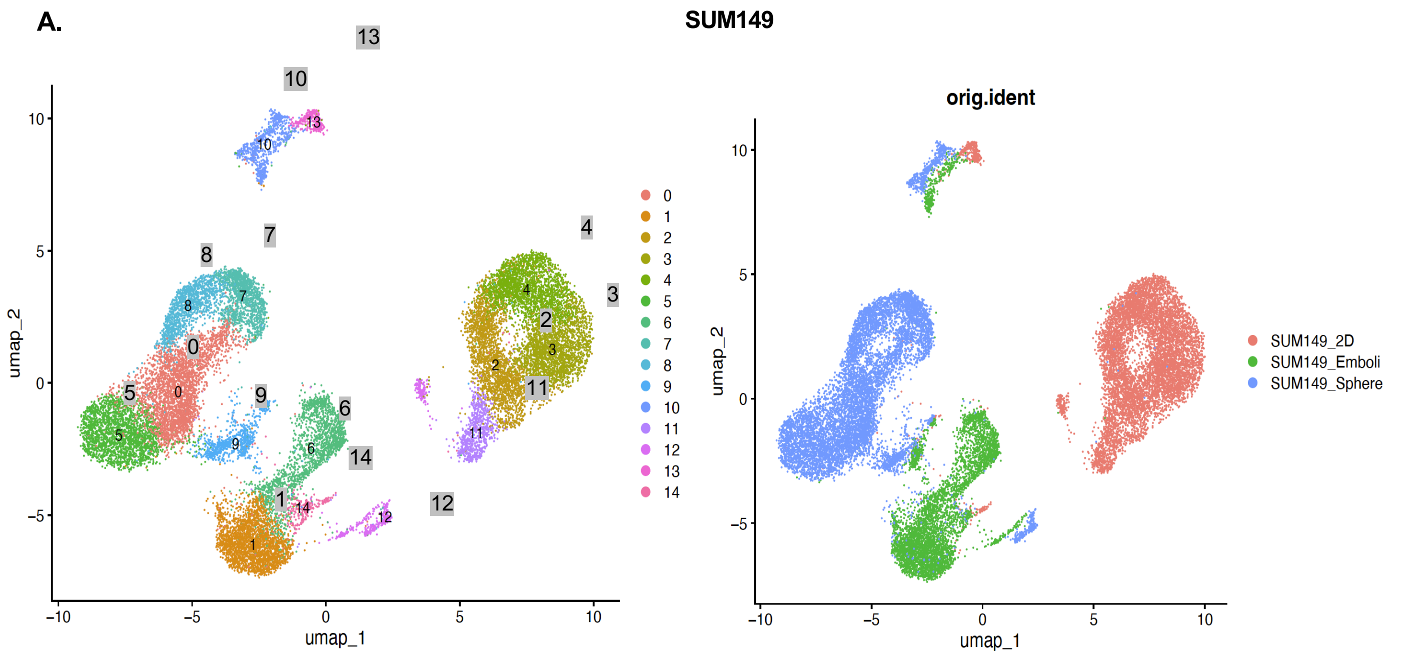


| group | 0 | 1 | 2 | 3 | 4 | 5 | 6 | 7 | 8 | 9 | 10 | 11 | 12 | 13 | 14 |
|---|---|---|---|---|---|---|---|---|---|---|---|---|---|---|---|
| SUM149_2D | 7 | 4 | 2408 | 2309 | 2253 | 2 | 18 | 4 | 2 | 33 | 15 | 725 | 153 | 277 | 64 |
| SUM149_Emboli | 6 | 2486 | 0 | 0 | 2 | 8 | 1818 | 3 | 3 | 259 | 294 | 1 | 103 | 17 | 164 |
| SUM149_Sphere | 3022 | 36 | 1 | 2 | 1 | 2225 | 33 | 1269 | 1269 | 468 | 444 | 3 | 199 | 7 | 13 |

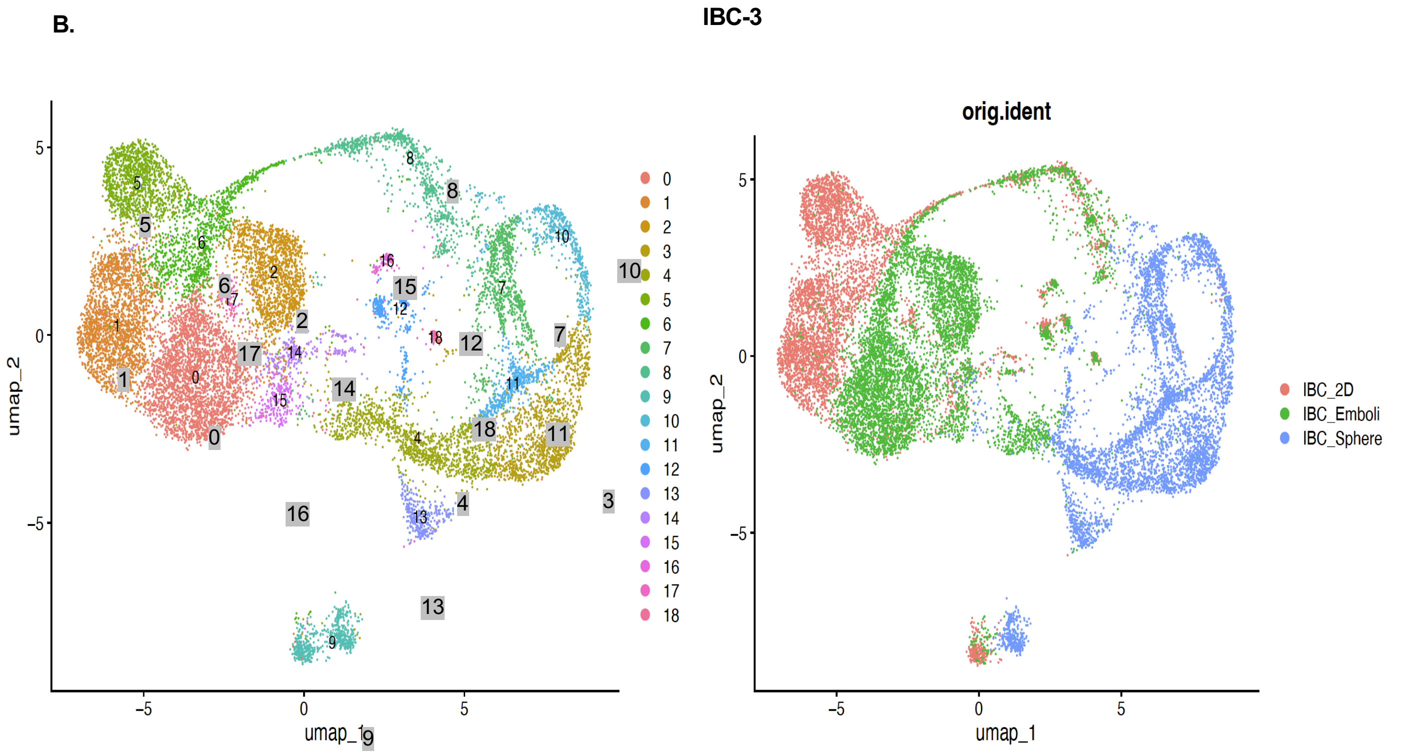


| group | 0 | 1 | 2 | 3 | 4 | 5 | 6 | 7 | 8 | 9 | 10 | 11 | 12 | 13 | 14 | 15 | 16 | 17 | 18 |
|---|---|---|---|---|---|---|---|---|---|---|---|---|---|---|---|---|---|---|---|
| IBC_2D | 115 | 2135 | 48 | 3 | 1 | 1251 | 563 | 9 | 294 | 209 | 7 | 0 | 113 | 0 | 104 | 64 | 51 | 50 | 33 |
| IBC_Emboli | 2745 | 36 | 1507 | 8 | 376 | 8 | 525 | 24 | 486 | 57 | 14 | 2 | 126 | 12 | 137 | 188 | 47 | 35 | 35 |
| IBC_Sphere | 5 | 0 | 1 | 1431 | 883 | 0 | 2 | 887 | 21 | 300 | 501 | 519 | 108 | 303 | 24 | 6 | 0 | 1 | 0 |

**Figure S2. Single cell mRNA sequencing of SUM149 and IBC-3 cells from 2D, SphC, and EmC.** UMAP clusters of scRNA-Seq data derived from (A) SUM149 and (B) IBC-3 cells after 3 days in the indicated culture conditions, along with tables showing the number of cells per cluster and condition. Data represent the combination of two biological replicates each.

**Figure S3**

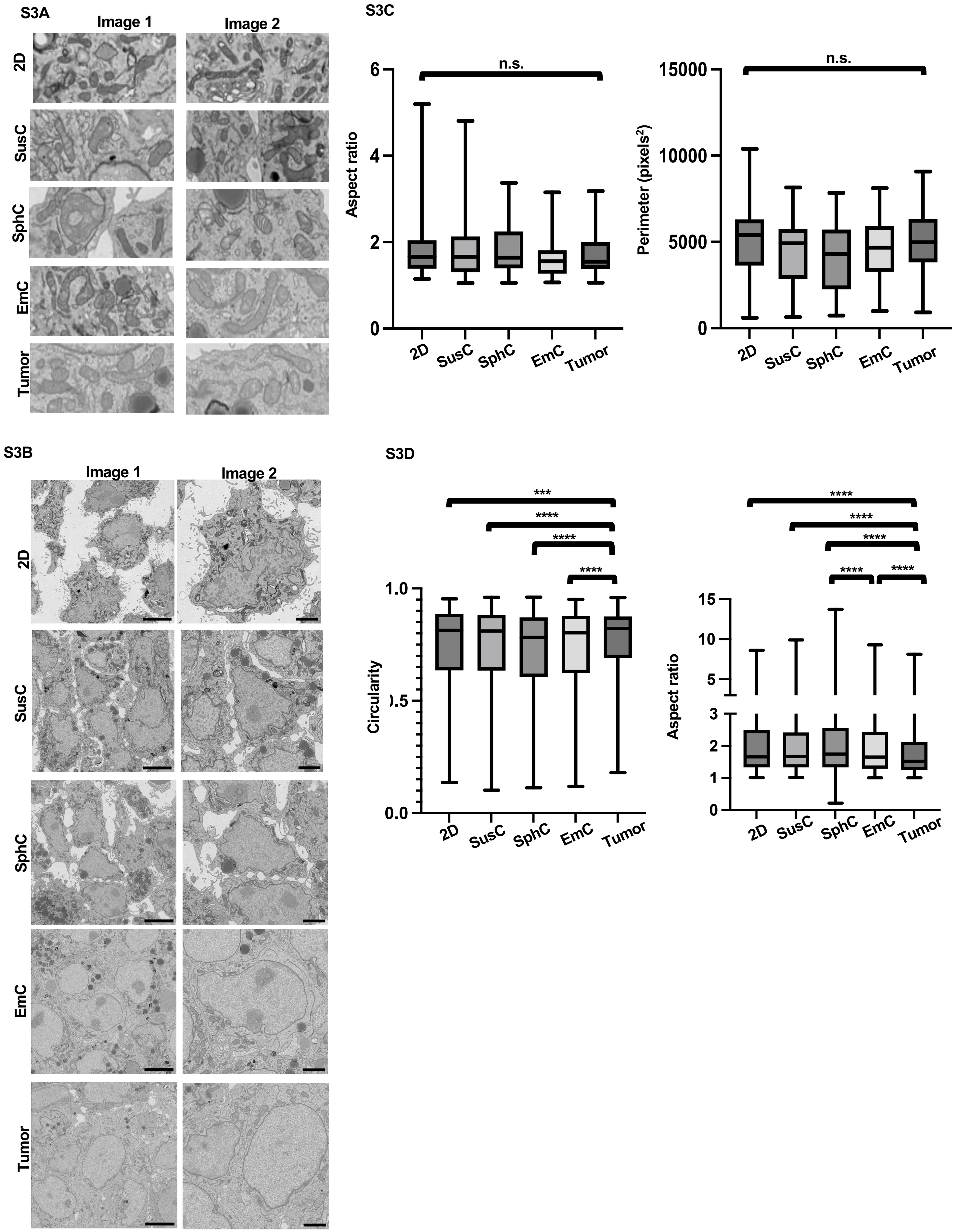
S3A
Image 1
Image 2
2D
SusC
SphC
EmC
Tumor
S3C
n.s.
Aspect ratio
6
4
2
0
2D
SusC
SphC
EmC
Tumor
n.s.
Perimeter (pixels$^2$)
15000
10000
5000
0
2D
SusC
SphC
EmC
Tumor
S3B
Image 1
Image 2
2D
SusC
SphC
EmC
Tumor
S3D
***
****
****
****
Circularity
1.0
0.0
2D
SusC
SphC
EmC
Tumor
****
****
****
****
****
Aspect ratio
15
10
5
3
2
1
0
2D
SusC
SphC
EmC
Tumor

**Figure S3**

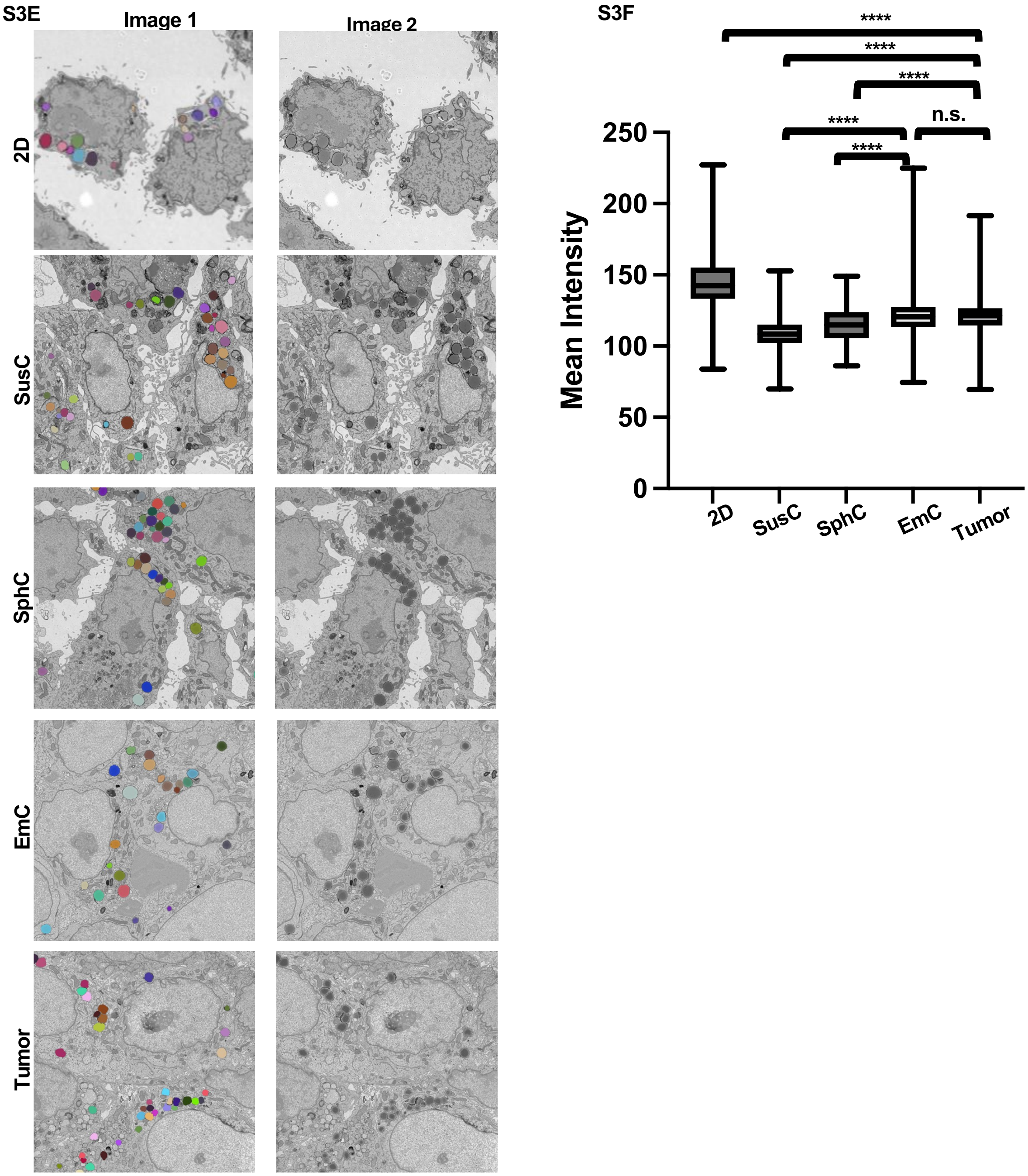


**Figure S3. Ultrastructural analysis of SUM149 mitochondria, nuclei and lipid droplets.**
**A,** Scanning electron microscope images showing mitochondria under indicated conditions. Images taken in FIJI at 25% zoom with 5 µm x 2 µm box. Pixel size = 5 nm. **B,** Scanning electron microscope images of nuclei as in A. Scale bar = 2 µm. **C,** Morphometric data of nuclei in cells cultured as indicated (n = 50-68, mean). **D,** Morphometric data of mitochondria in cells cultured as indicated (n = 1451-1995). **E,** Scanning electron microscope images of lipid droplets LDs (as identified in the image on the left) in cells cultured as indicated. Pixel size=5 nm. **F,** Morphometric data of lipid droplets in cells cultured as indicated (n = 131-654).
***$P$<0.0001, ****$P$<0.00001, n.s. not significant.

### Figure S4

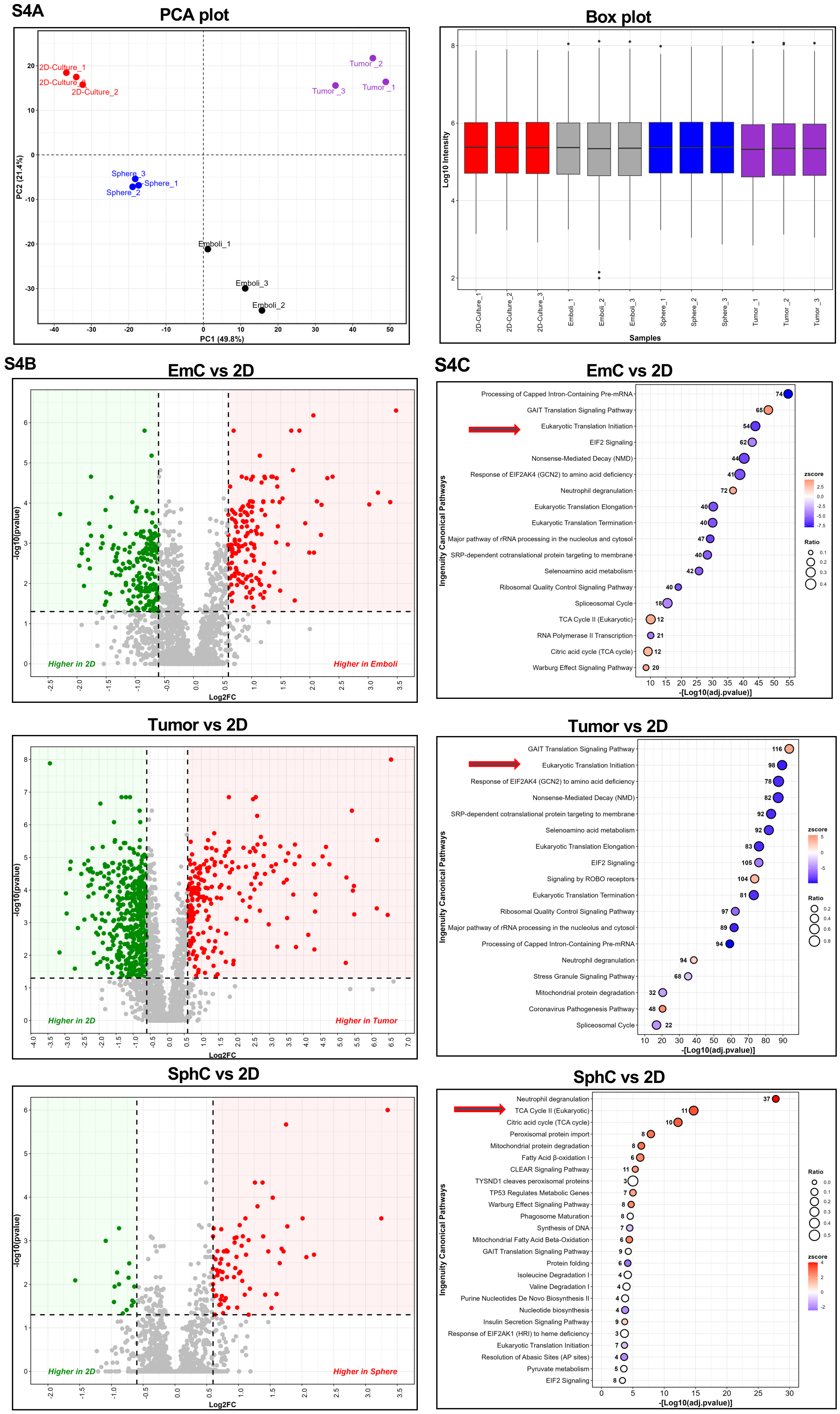
S4A
PCA plot
2D-Culture_1
2D-Culture_3
2D-Culture_2
Tumor_2
Tumor_3
Tumor_1
Sphere_3
Sphere_1
Sphere_2
Emboli_1
Emboli_3
Emboli_2
PC1 (49.8%)
PC2 (21.4%)
Box plot
Log10 Intensity
Samples
2D-Culture_1
2D-Culture_2
2D-Culture_3
Emboli_1
Emboli_2
Emboli_3
Sphere_1
Sphere_2
Sphere_3
Tumor_1
Tumor_2
Tumor_3
S4B
EmC vs 2D
-log10(pvalue)
Log2FC
Higher in 2D
Higher in Emboli
S4C
EmC vs 2D
Processing of Capped Intron-Containing Pre-mRNA
GAIT Translation Signaling Pathway
Eukaryotic Translation Initiation
EIF2 Signaling
Nonsense-Mediated Decay (NMD)
Response of EIF2AK4 (GCN2) to amino acid deficiency
Neutrophil degranulation
Eukaryotic Translation Elongation
Eukaryotic Translation Termination
Major pathway of rRNA processing in the nucleolus and cytosol
SRP-dependent cotranslational protein targeting to membrane
Selenoamino acid metabolism
Ribosomal Quality Control Signaling Pathway
Spliceosomal Cycle
TCA Cycle II (Eukaryotic)
RNA Polymerase II Transcription
Citric acid cycle (TCA cycle)
Warburg Effect Signaling Pathway
Ingenuity Canonical Pathways
-[Log10(adj.pvalue)]
zscore
Ratio
Tumor vs 2D
-log10(pvalue)
Log2FC
Higher in 2D
Higher in Tumor
Tumor vs 2D
GAIT Translation Signaling Pathway
Eukaryotic Translation Initiation
Response of EIF2AK4 (GCN2) to amino acid deficiency
Nonsense-Mediated Decay (NMD)
SRP-dependent cotranslational protein targeting to membrane
Selenoamino acid metabolism
Eukaryotic Translation Elongation
EIF2 Signaling
Signaling by ROBO receptors
Eukaryotic Translation Termination
Ribosomal Quality Control Signaling Pathway
Major pathway of rRNA processing in the nucleolus and cytosol
Processing of Capped Intron-Containing Pre-mRNA
Neutrophil degranulation
Stress Granule Signaling Pathway
Mitochondrial protein degradation
Coronavirus Pathogenesis Pathway
Spliceosomal Cycle
Ingenuity Canonical Pathways
-[Log10(adj.pvalue)]
zscore
Ratio
SphC vs 2D
-log10(pvalue)
Log2FC
Higher in 2D
Higher in Sphere
SphC vs 2D
Neutrophil degranulation
TCA Cycle II (Eukaryotic)
Citric acid cycle (TCA cycle)
Peroxisomal protein import
Mitochondrial protein degradation
Fatty Acid β-oxidation I
CLEAR Signaling Pathway
TYSND1 cleaves peroxisomal proteins
TP53 Regulates Metabolic Genes
Warburg Effect Signaling Pathway
Phagosome Maturation
Synthesis of DNA
Mitochondrial Fatty Acid Beta-Oxidation
GAIT Translation Signaling Pathway
Protein folding
Isoleucine Degradation I
Valine Degradation I
Purine Nucleotides De Novo Biosynthesis II
Nucleotide biosynthesis
Insulin Secretion Signaling Pathway
Response of EIF2AK1 (HRI) to heme deficiency
Eukaryotic Translation Initiation
Resolution of Abasic Sites (AP sites)
Pyruvate metabolism
EIF2 Signaling
Ingenuity Canonical Pathways
-[Log10(adj.pvalue)]
Ratio
zscore

Figure S4

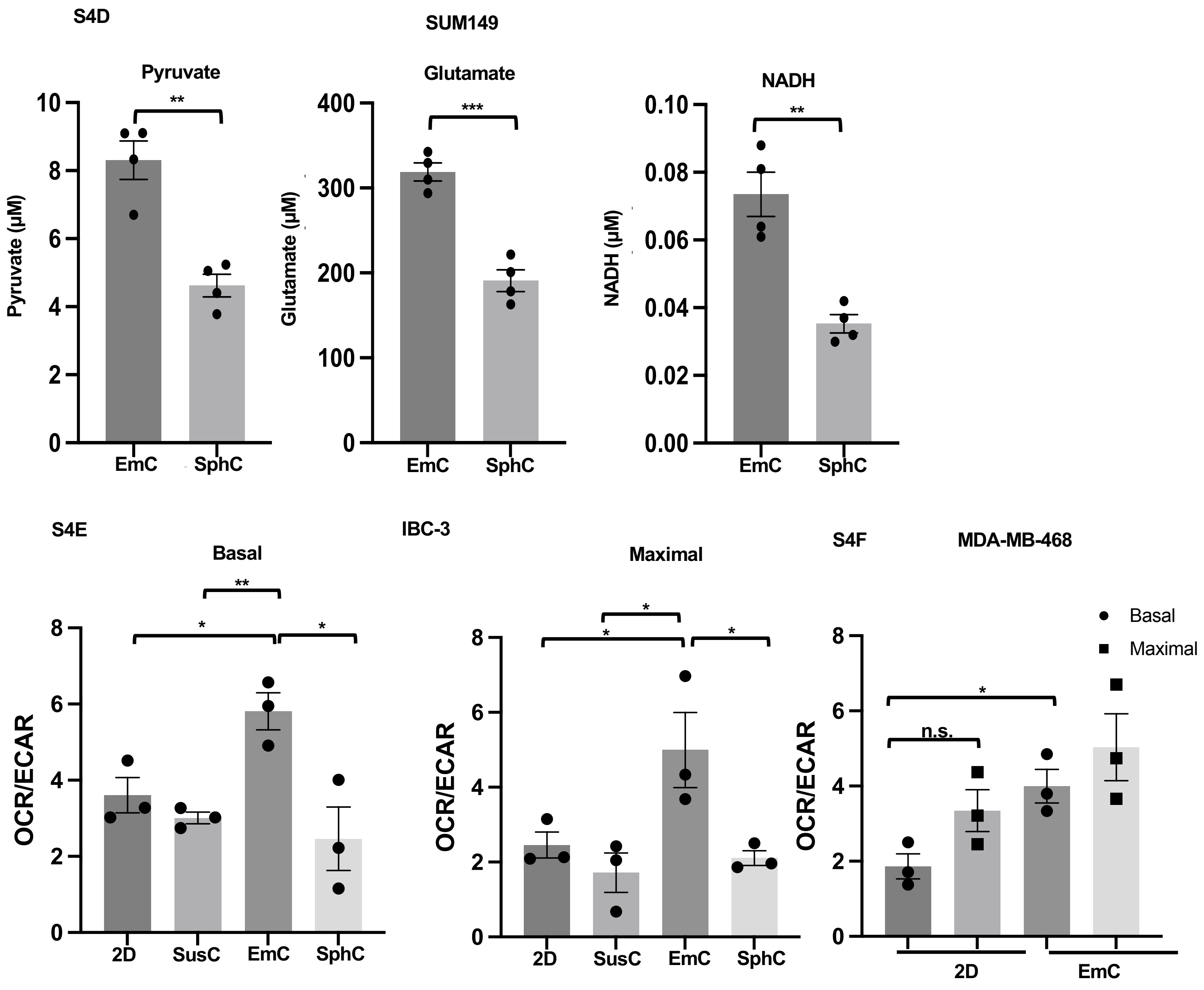


**Figure S4. Proteomic and metabolic analysis reveals differential cell adaptations to 3D culture.** **A,** PCA and box plots of proteomic data (Supplemental File 5) from SUM149 cells in different culture conditions and xenograft tumor tissue (n = 3). **B,** Volcano plots of the data in panel A. **C,** Ingenuity Pathway Analysis of protein expression data as in panels A-B. Arrows point to pathways discussed in the Results. **D,** Intracellular pyruvate, glutamate and NADH concentrations in SUM149 cells cultured for 3 days in SphC and EmC (n = 4). **E,** OCR/ECAR ratio (basal and maximal) in IBC-3 cells after 3 days under the indicated culture conditions. Data are from 3 different experiments with 3 time points and 3-4 technical replicates each. **F,** OCR/ECAR ratio (basal and maximal) in MDA-MB-468 cells cultured as 2D and EmC. Data are from 3 different experiments with 3 time points and 3-4 technical replicates each. Data are mean ± SEM,*P<0.05, **P<0.01, ***P<0.001, n.s., not significant.

Figure S5

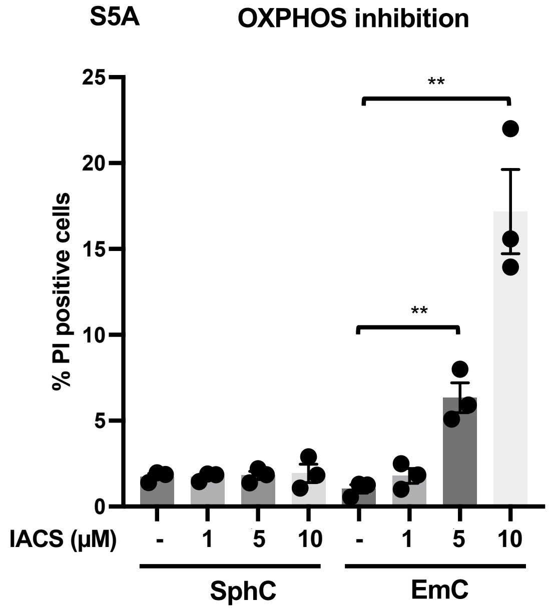


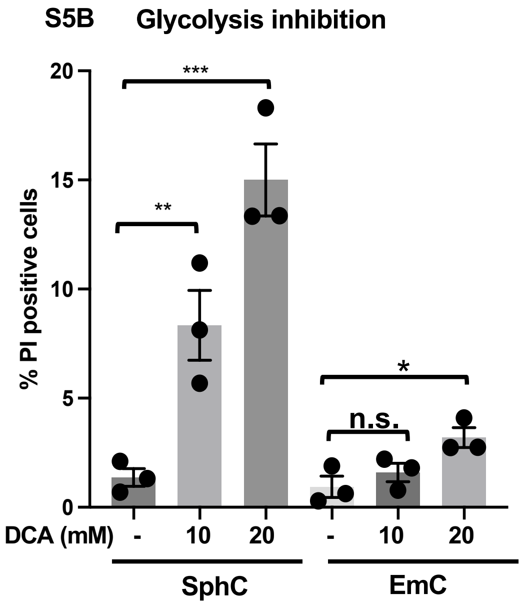


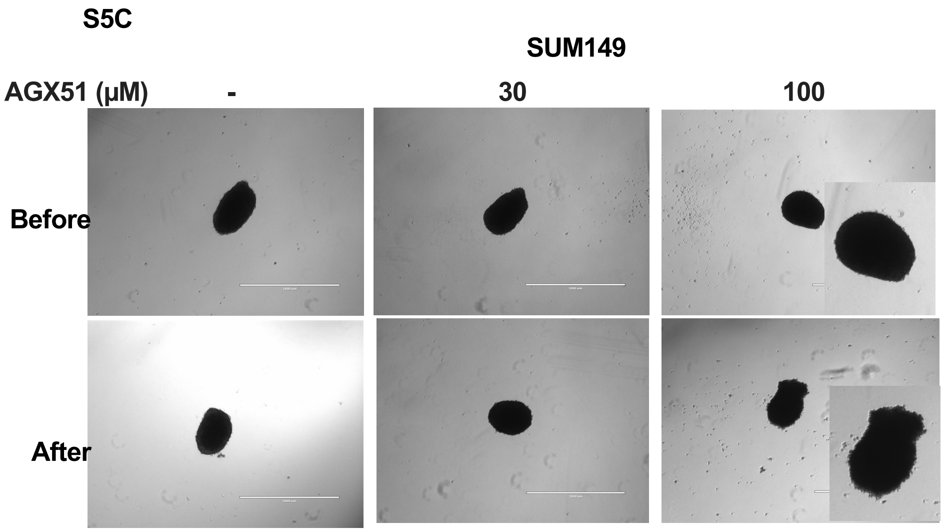


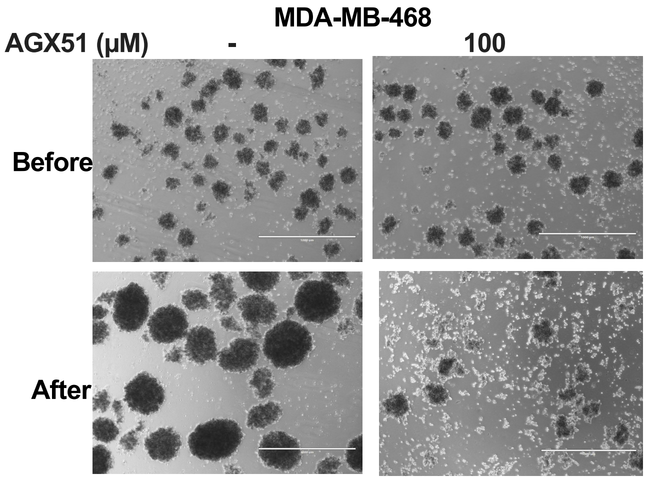


**Figure S5. Cells in EmC are sensitive to OXPHOS and ID1/ID3 inhibitors.**

**A,** Percent PI positive cells in EmC treated with different doses of IACS-010579 for 72 h. **B,** Percent PI positive cells in SphC treated with different doses of DCA for 72 h. **C,** Bright filed images of SUM149 and MDA-MB-468 cells in EmC before and after treatment with ID1/ID3 inhibitor, AGX51. Scale bar=1 mm. Inset shows high magnification to visualize fraying periphery of the embolus in the presence of AGX51 of SUM159 cells. Data are mean ± SEM,**P*<0.05, ***P*<0.01, ****P*<0.001, n.s., not significant.
